# Nutrient flux governs osteogenic fate commitment through the SLC3A1-cystine Axis

**DOI:** 10.64898/2026.07.25.740706

**Authors:** Yangshuai Gao, Qixuan Lin, Qing Xu, Shaokang Chen, Lei Jiang, Yiming Hu, Junrong Cai, Aaron W. James, Zhongmin Zhang, Jiajia Xu

## Abstract

Skeletal and mesenchymal cells have limited bone-forming potential due to their rarity, tendency for senescence and/or unstable osteogenic lineage commitment. We previously identified an osteopotent CXCR4⁺ stem cell population with unclear mechanism for its differentiation potential. Here, we found that the cystine transporter SLC3A1 was selectively enriched in CXCR4^+^ stem cells, uncovering a role for amino acid transport in regulating osteogenic fate commitment. Enforced SLC3A1 expression reprogrammed mesenchymal cells toward a stable osteogenic state while suppressing adipogenic differentiation. Mechanistically, SLC3A1-mediated cystine flux established a glutathione-dependent metabolic program that preserved mitochondrial fitness and restrained stem cell senescence. SLC3A1 also stabilized IGF-1 through suppression of ZMYND8-mediated ubiquitination, uncovering ZMYND8 as a previously unrecognized E3 ligase regulating osteogenic commitment. Cystine supplementation phenocopied the effects of SLC3A1 activation, promoting osteogenic differentiation and skeletal regeneration without genetic manipulation. *In vivo*, cystine administration accelerated bone regeneration and attenuated ovariectomy-induced bone loss, while analyses of a human osteoporosis cohort, osteoporotic specimens, and single-cell transcriptomic datasets revealed coordinated suppression of the SLC3A1-cystine-IGF-1 pathway in osteoporotic mesenchymal cells. Taken together, these findings establish SLC3A1-mediated cystine transport as a programmable metabolic determinant of osteogenic fate commitment and identify cystine metabolism as a therapeutically targetable axis for skeletal regeneration and osteoporosis.

## Introduction

Mesenchymal stem/stromal cells (MSCs) hold substantial promise for skeletal regeneration because of their accessibility, multilineage differentiation capacity, and regenerative potential^1–3^. However, progressive loss of osteogenic competence and functional heterogeneity remain major barriers limiting the efficacy and reproducibility of stem cell-mediated bone repair^4–8^. In particular, prolonged *ex vivo* expansion frequently leads to impaired osteogenic differentiation, cellular senescence, and regenerative decline, thereby compromising the translational potential of stem cell-based skeletal therapies^4,9–11^. Despite extensive efforts to improve regenerative efficacy, the mechanisms that establish and maintain stable osteogenic fate remain incompletely understood.

Emerging evidence has established metabolism as a fundamental regulator of stem cell fate transitions^12–15^. Beyond bioenergetic support, metabolic pathways actively coordinate epigenetic remodeling, redox adaptation, and lineage specification^12,14,16,17^. In MSCs, amino acid metabolism, glutathione-dependent redox homeostasis, and mitochondrial activity critically regulate osteogenic differentiation, cellular fitness, and skeletal maintenance^18–23^. Conversely, oxidative and replicative stress accumulated during *in vitro* expansion promote stem cell dysfunction and compromise regenerative capacity^13,24,25^. These observations suggest that metabolic adaptation may represent a central mechanism governing osteogenic competence. However, the specific metabolic programs that stabilize osteogenic fate and preserve regenerative function remain poorly defined.

We previously identified a rare C-X-C motif chemokine receptor 4^+^ (CXCR4⁺) mesenchymal cell population associated with enhanced osteogenic potential and robust skeletal regeneration^26^. However, these cells are extremely scarce, exhibit substantial inter-donor variability, and lose regenerative competence during *ex vivo* expansion. More importantly, the molecular mechanisms that establish and maintain this osteogenic state remain undefined. We therefore reasoned that defining the metabolic adaptations underlying this phenotype could reveal strategies to stabilize osteogenic fate and overcome functional heterogeneity.

In this study, we identified a metabolically specialized osteogenic state characterized by activation of a solute carrier family 3 member 1 (SLC3A1)-cystine signaling axis. Mechanistically, SLC3A1 expression was driven by SMAD3 downstream of transforming growth factor-beta (TGF-β) signaling and promoted intracellular cystine utilization to establish a redox-adaptive metabolic program that preserves mitochondrial fitness and suppresses cellular senescence. We further identified the previously unrecognized E3 ligase zinc finger MYND-type containing 8 (ZMYND8) as a key regulator of insulin-like growth factor 1 (IGF-1) stability and demonstrated that SLC3A1-cystine signaling sustained osteogenic commitment by preventing ZMYND8-mediated IGF-1 degradation. Notably, either enforced SLC3A1 expression or exogenous cystine supplementation was sufficient to reprogram bulk MSCs toward a CXCR4⁺-like osteogenic state, thereby enhancing bone formation and attenuating osteoporotic bone loss in both experimental models and human specimens. Collectively, these findings establish the SLC3A1-cystine metabolic axis as a central regulator of osteogenic fate and uncover a metabolically driven strategy for functional cell-state reprogramming to enhance skeletal regeneration.

## Results

### CXCR4 marks an osteogenic cell state but does not directly regulate osteogenic commitment

To identify molecular mechanisms underlying osteogenic competence, we first isolated MSCs based on surface CXCR4 expression (**Fig. 1A**). Consistent with our previous findings, CXCR4^+^ cells exhibited significantly enhanced osteogenic capacity compared with CXCR4^-^ cells, as evidenced by elevated mineralization (**Fig. 1B**) and increased expression of *CXCR4* and osteogenic markers such as RUNX family transcription factor 2 (*RUNX2*), osterix (*SP7*), and osteocalcin (*BGLAP*) (**Fig. 1C-F**). These observations confirmed that surface CXCR4 expression identifies a distinct osteogenic cell state associated with enhanced regenerative potential.

**Figure 1.**
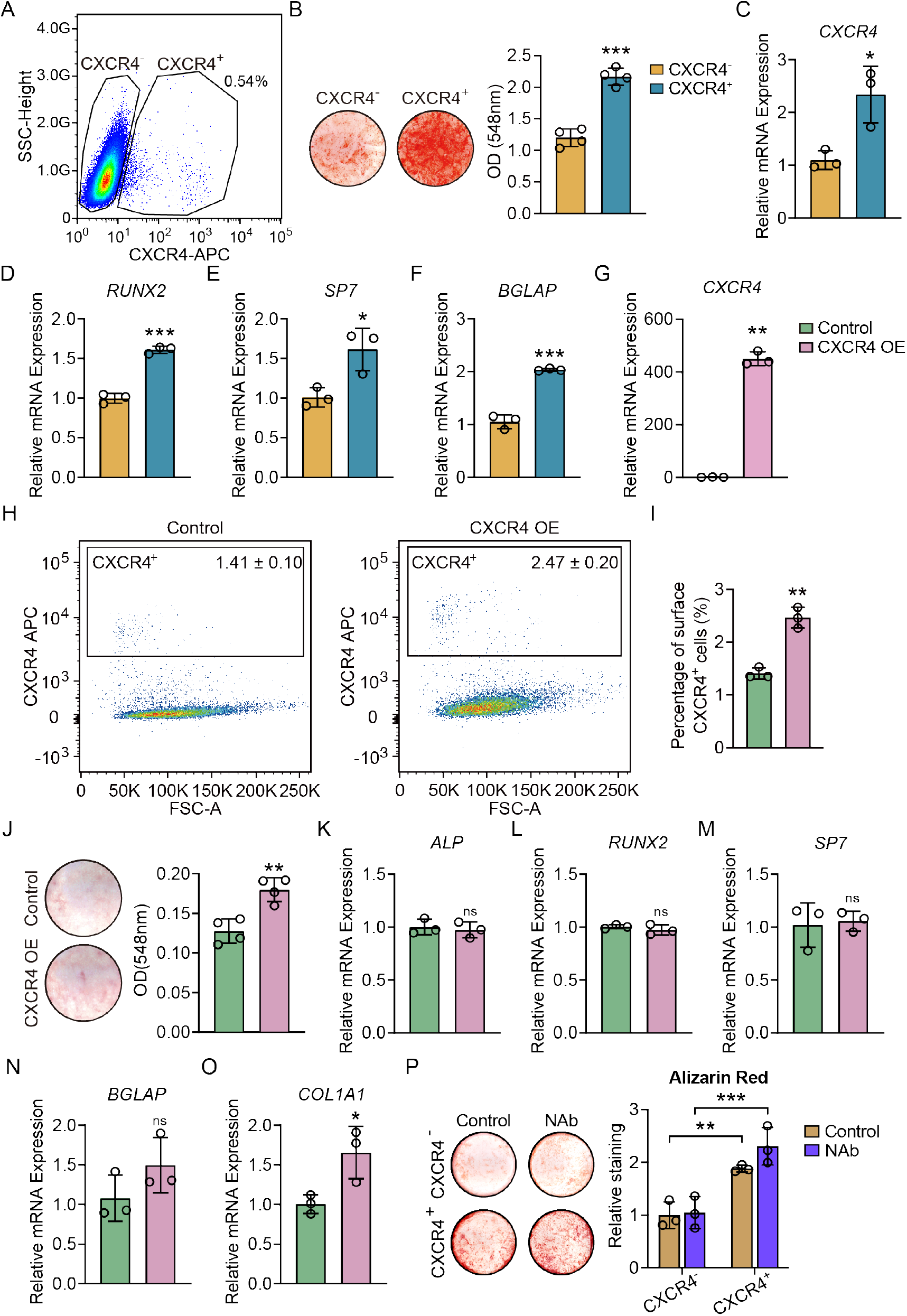
CXCR4 marks an osteogenic cell state but does not directly regulate osteogenic commitment. **(A)** Representative flow cytometry plot showing the gating strategy used to isolate CXCR4^+^ and CXCR4^-^ mesenchymal cells based on surface CXCR4 expression. **(B)** Alizarin Red S staining of CXCR4^+^ and CXCR4^-^ cells after 6 days of osteogenic induction, showing enhanced mineral deposition in CXCR4^+^ cells. **(C-F)** qPCR analysis of *CXCR4* **(C)**, *RUNX2* **(D)**, *SP7* **(E)**, and *BGLAP* **(F)** expression in CXCR4^-^ and CXCR4^+^ cells. **(G)** qPCR analysis confirming CXCR4 overexpression efficiency. **(H,I)** Representative flow cytometry plots and quantification showing modest increase in the percentage of CXCR4^+^ cells following CXCR4 overexpression. **(J)** Alizarin Red S staining showed a slight increase in mineralized matrix formation following CXCR4 overexpression. **(K-O)** qPCR analysis of canonical osteogenic markers *ALP* **(K)**, *RUNX2* **(L)**, *SP7* **(M)**, *BGLAP* **(N)**, and *COL1A1* **(O)** in control and CXCR4-overexpressing cells. ns, not significant. **(P)** Alizarin Red S staining showing that blocking surface CXCR4 with a neutralizing antibody did not impair the osteogenic capacity of CXCR4^+^ cells. Data are presented as mean ± SD. \**P*<0.05; \*\**P*<0.01; \*\*\**P*<0.001.

Because CXCR4⁺ cells represent only a small fraction of the total mesenchymal cell population, we next asked whether CXCR4 itself functionally contributes to osteogenic commitment. Forced CXCR4 expression in bulk MSCs resulted in only a modest increase in surface CXCR4 positivity (**Fig. 1G-I**) and produced minimal effects on osteogenic differentiation, as assessed by Alizarin Red S staining (**Fig. 1J**) and expression of alkaline phosphatase (*ALP*), *RUNX2*, *SP7*, and *BGLAP* (**Fig. 1K-O**).

To further evaluate the functional contribution of CXCR4 signaling, we blocked surface CXCR4 using a neutralizing antibody. Unexpectedly, inhibition of CXCR4 signaling failed to impair osteogenic differentiation of CXCR4^+^ cells, as assessed by Alizarin Red S staining (**Fig. 1P**). Taken together, these findings indicate that CXCR4 serves primarily as a phenotypic marker of osteogenic competence rather than a functional regulator of osteogenic commitment. These results further suggest that the enhanced regenerative properties of CXCR4^+^ cells are likely governed by downstream molecular programs independent of CXCR4 receptor signaling.

### Transcriptional landscape of CXCR4^+^ mesenchymal cells reveals osteogenic bias and highlights SLC3A1 as a candidate regulator

To elucidate the molecular mechanisms underlying the enhanced osteogenic competence, we performed bulk RNA sequencing on CXCR4^+^ and CXCR4^-^ MSCs. Principal component analysis (PCA) revealed a clear transcriptional separation between the two groups (**Supplementary Fig. 1A**), further supported by hierarchical clustering and volcano plot analysis, which highlighted substantial differences in gene expression profiles (**Supplementary Fig. 1B,C**). Among the differentially expressed genes, *PRSS3*, *KRT14*, *TSPAN8*, *PCDH10*, *LYPD1*, *LHX9*, *CNTNAP3B*, and *SLC3A1* were the top eight genes significantly upregulated in CXCR4^+^ cells (**Supplementary Fig. 1D**). Functional enrichment analyses revealed a strong osteogenic signature in CXCR4^+^ cells. Gene Ontology (GO) analysis showed that CXCR4^+^ cells were enriched for biological processes related to skeletal system development, extracellular matrix, and ossification (**Supplementary Fig. 1E**). KEGG pathway analysis revealed activation of multiple signaling pathways including TGF-beta signaling, MAPK signaling, and cytokine-cytokine receptor interaction (**Supplementary Fig. 1F**). Differential expression analysis of lineage-associated genes demonstrated that osteogenic genes were significantly enriched in CXCR4^+^ cells, whereas adipogenic markers were more abundant in the CXCR4^-^ population (**Supplementary Fig. 1G**), consistent with the functional osteogenic bias observed *in vitro*. Gene Set Enrichment Analysis (GSEA) further confirmed upregulation of ossification-related pathways together with suppression of lipid metabolism programs in the CXCR4⁺ subset (**Supplementary Fig. 1H,I**).

To identify candidate regulators responsible for this osteogenic state, we validated the top differentially expressed genes by qPCR. Among the candidates examined, SLC3A1 displayed the most robust and reproducible enrichment in CXCR4^+^ cells (**Supplementary Fig. 1J and Supplementary Fig. 2**). Notably, SLC3A1 expression progressively increased throughout osteogenic induction, suggesting a potential role in establishing and maintaining osteogenic commitment (**Supplementary Fig. 1K**). Therefore, these data identify SLC3A1 as a prominent molecular feature of osteogenic-competent mesenchymal cells and nominate it as a candidate regulator of osteogenic fate.

### SLC3A1 reprograms stem cells toward an osteogenic precursor identity

We next conducted gain- and loss-of-function experiments to determine whether SLC3A1 contributes to osteogenic fate regulation in MSCs. Overexpression of SLC3A1 markedly enhanced osteogenesis, as demonstrated by increased Alizarin Red S staining and elevated the expression of canonical osteogenic markers including *RUNX2*, *ALP*, and *collagen type I alpha 1* (*COL1A1*) (**Fig. 2A,B,I**). Conversely, silencing *SLC3A1* with siRNA or pharmacologically inhibiting its activity with Sulfasalazine^27^ significantly impaired osteogenic differentiation, resulting in reduced mineral deposition and diminished expression of osteogenic marker genes (**Fig. 2C-F,I**). These findings establish SLC3A1 as a critical regulator of osteogenic commitment.

**Figure 2.**
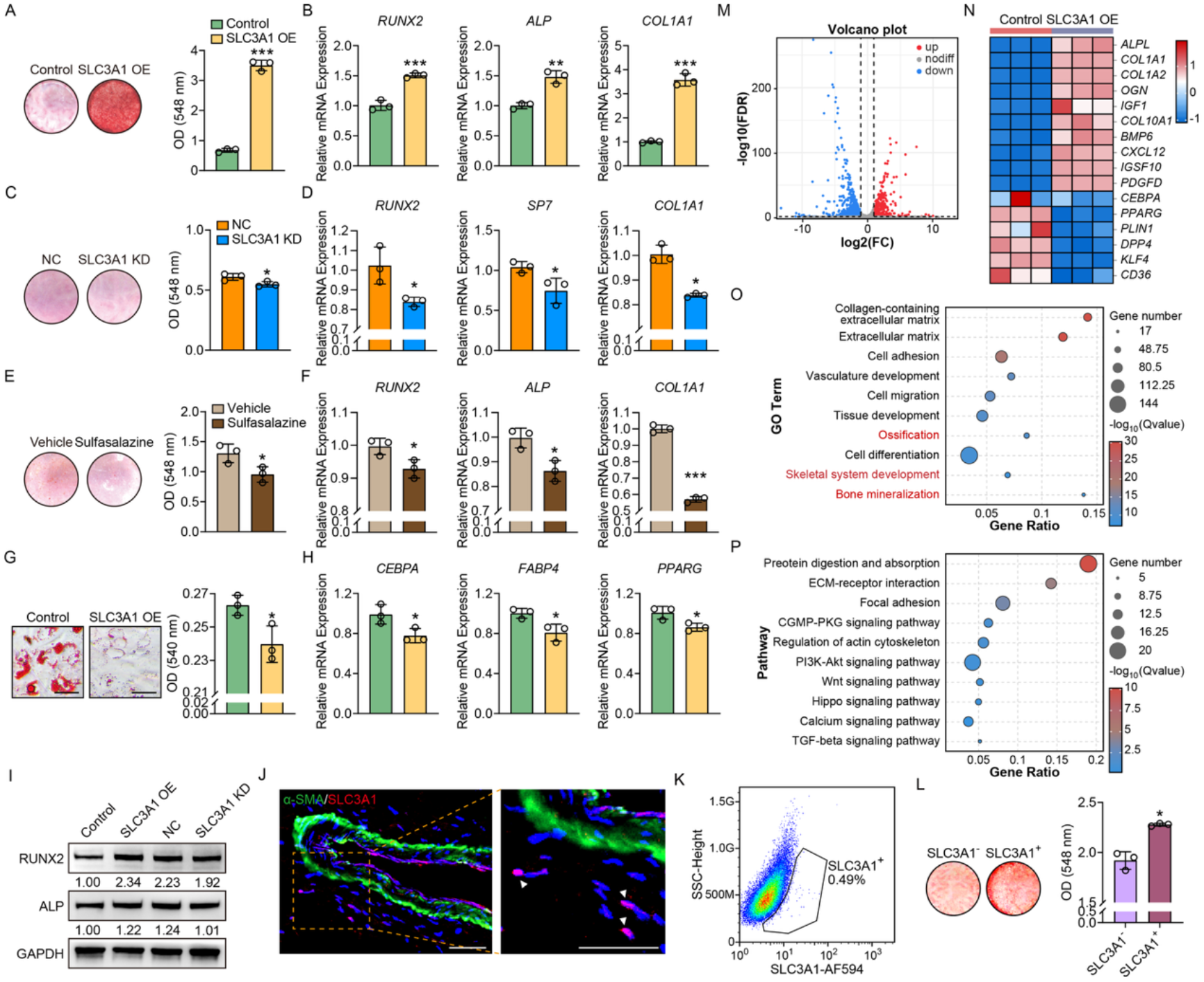
SLC3A1 promotes osteogenic commitment and establishes an osteogenic cell state in adipose-derived MSCs. **(A)** Representative images of Alizarin Red S staining showing enhanced mineralized matrix deposition following SLC3A1 overexpression. **(B)** qPCR analysis of canonical osteogenic markers (*RUNX2*, *ALP*, *COL1A1*) in SLC3A1-overexpressing MSCs. **(C,D)** Alizarin Red S staining and quantification showing impaired osteogenesis upon SLC3A1 knockdown by siRNA. **(E,F)** Alizarin Red S staining and osteogenic marker gene expression after pharmacologic inhibition of SLC3A1 with sulfasalazine (0.5 mM). **(G)** Oil Red O staining showing reduced lipid droplet formation in SLC3A1-overexpressing MSCs. **(H)** qPCR analysis of adipocyte-associated genes (*CEBPA*, *FABP4*, *PPARG*) in control and SLC3A1-overexpressing MSCs. **(I)** Western blot demonstrated the expression of RUNX2 and ALP across SLC3A1 gain- and loss-of-function conditions. **(J)** Immunofluorescence staining of SLC3A1 in the vascular adventitia. α-SMA was used to delineate vascular smooth muscle regions, confirming perivascular localization of SLC3A1^+^ cells (indicated by white arrowheads). Nuclei were counterstained with DAPI (blue). **(K)** Flow cytometry quantification of SLC3A1^+^ cell frequencies in adipose-derived MSC populations. **(L)** Alizarin Red S staining and quantification comparing osteogenic capacity of sorted SLC3A1^-^ and SLC3A1^+^ MSCs. **(M-P)** Transcriptomic reprogramming in SLC3A1-overexpressing MSCs favors osteogenic over adipogenic fate. **(M)** Volcano plot of DEGs (red: upregulated in SLC3A1 OE group, blue: downregulated in SLC3A1-overexpressing group; FDR < 0.05, |log_2_FC| > 1). **(N)** Heatmap of representative osteogenic and adipogenic lineage genes, showing enhanced osteogenic and suppressed adipogenic signatures in SLC3A1-overexpressing cells. **(O,P)** GO and KEGG pathway enrichment analysis of upregulated genes, highlighting terms related to ossification, skeletal system development, and Wnt signaling. OE, overexpression; KD, knockdown; NC, negative control. Scale bars: 50 μm. Data are presented as mean ± SD. \**P*<0.05; \*\**P*<0.01; \*\*\**P*<0.001.

To determine whether SLC3A1 influences lineage specification beyond osteogenesis, we next evaluated adipogenic differentiation. Oil Red O staining revealed that SLC3A1 overexpression suppressed lipid droplet accumulation, accompanied by decreased transcription of adipocyte-associated genes *CCAAT enhancer binding protein alpha* (*CEBPA*), *fatty acid binding protein 4* (*FABP4*), and *peroxisome proliferator activated receptor gamma* (*PPARG*) (**Fig. 2G,H**). Immunofluorescence confirmed the presence of SLC3A1^+^ cells within perivascular regions (**Fig. 2J**), and flow cytometry revealed a frequency comparable to that of CXCR4^+^ cells (**Fig. 2K**). Notably, SLC3A1^+^ mesenchymal cells displayed superior osteogenic capacity relative to SLC3A1^-^ cells (**Fig. 2L**), suggesting that SLC3A1 marks and promotes an osteo-primed phenotype. Thus, these results indicate that SLC3A1 promotes a shift toward osteogenic fate while simultaneously restraining adipogenic differentiation. The phenotypic resemblance between SLC3A1^+^ cells and the CXCR4^+^ subset further supports SLC3A1 as a key molecular regulator of osteogenic fate commitment.

To define the transcriptional programs downstream of SLC3A1, we performed bulk RNA sequencing of MSCs overexpressing SLC3A1. PCA revealed a distinct separation in the global transcriptional profiles between control and SLC3A1-overexpressing cells, indicating widespread transcriptomic reprogramming (**Supplementary Fig. 3A**). Differential gene expression analysis identified 378 genes significantly upregulated and 911 genes downregulated (>2-fold change, adjusted *P* value < 0.05; **Fig. 2M and Supplementary Fig. 3B**). Among the most highly induced transcripts were *NRGN*, *INSRR*, *SMOC2*, *COL10A1*, *IGF1*, and *IGFBP3*, many of which have been associated with skeletal development or stem cell activation (**Supplementary Fig. 3C**). Functional enrichment analyses revealed a strong osteogenic transcriptional program downstream of SLC3A1. GO and KEGG analyses identified enrichment of pathways associated with ossification, skeletal system development, bone mineralization, and Wnt signaling activation (**Fig. 2O,P**). Heatmap visualization and GSEA further confirmed increased expression of osteogenesis-associated genes, including *COL1A1*, *BMP6*, *ALPL*, *IGF1*, and *PDGFD*, together with enrichment of bone mineralization gene signatures in SLC3A1-overexpressing cells (**Fig. 2N and Supplementary Fig. 3D**). Conversely, genes associated with lipid storage and adipogenesis were downregulated (**Fig. 2N and Supplementary Fig. 3E**). Importantly, the global transcriptional changes induced by SLC3A1 closely resembled those observed in naturally osteogenic-competent CXCR4^+^ cells. Altogether, these findings establish SLC3A1 as a key regulator of an osteogenic cell state and demonstrate that activation of SLC3A1 is sufficient to confer transcriptional features associated with enhanced osteogenic competence.

### TGF-β/SMAD3 signaling activates *SLC3A1* transcription

Given the preferential expression of SLC3A1 in osteogenic-competent mesenchymal cells and its role in promoting osteogenic commitment, we next sought to elucidate the upstream regulatory mechanism responsible for *SLC3A1* transcriptional activation. Notably, overexpression of CXCR4 failed to alter *SLC3A1* mRNA levels (**Supplementary Fig. 4A**), further confirming that CXCR4 functions merely as a phenotypic marker rather than acting upstream in the signaling pathway that regulates SLC3A1 expression.

To identify signaling pathways responsible for SLC3A1 induction, we reanalyzed the transcriptomic profiles of CXCR4^+^ and CXCR4^-^ mesenchymal cell populations. KEGG and GSEA analyses revealed a significant enrichment of TGF-β signaling in the CXCR4^+^ cells (**Supplementary Fig. 1F and 4B**). We therefore investigated whether this pathway regulates SLC3A1 expression. qPCR analysis showed that stimulation with TGF-β1 significantly increased SLC3A1 mRNA levels (**Supplementary Fig. 4C**), while treatment with the selective TGF-β receptor inhibitor SB431542 effectively blocked this upregulation (**Fig. 3A**). Furthermore, knockdown of SMAD3, a key transcription factor downstream of TGF-β signaling, resulted in reduced SLC3A1 expression (**Supplementary Fig. 4D,E**).

**Figure 3.**
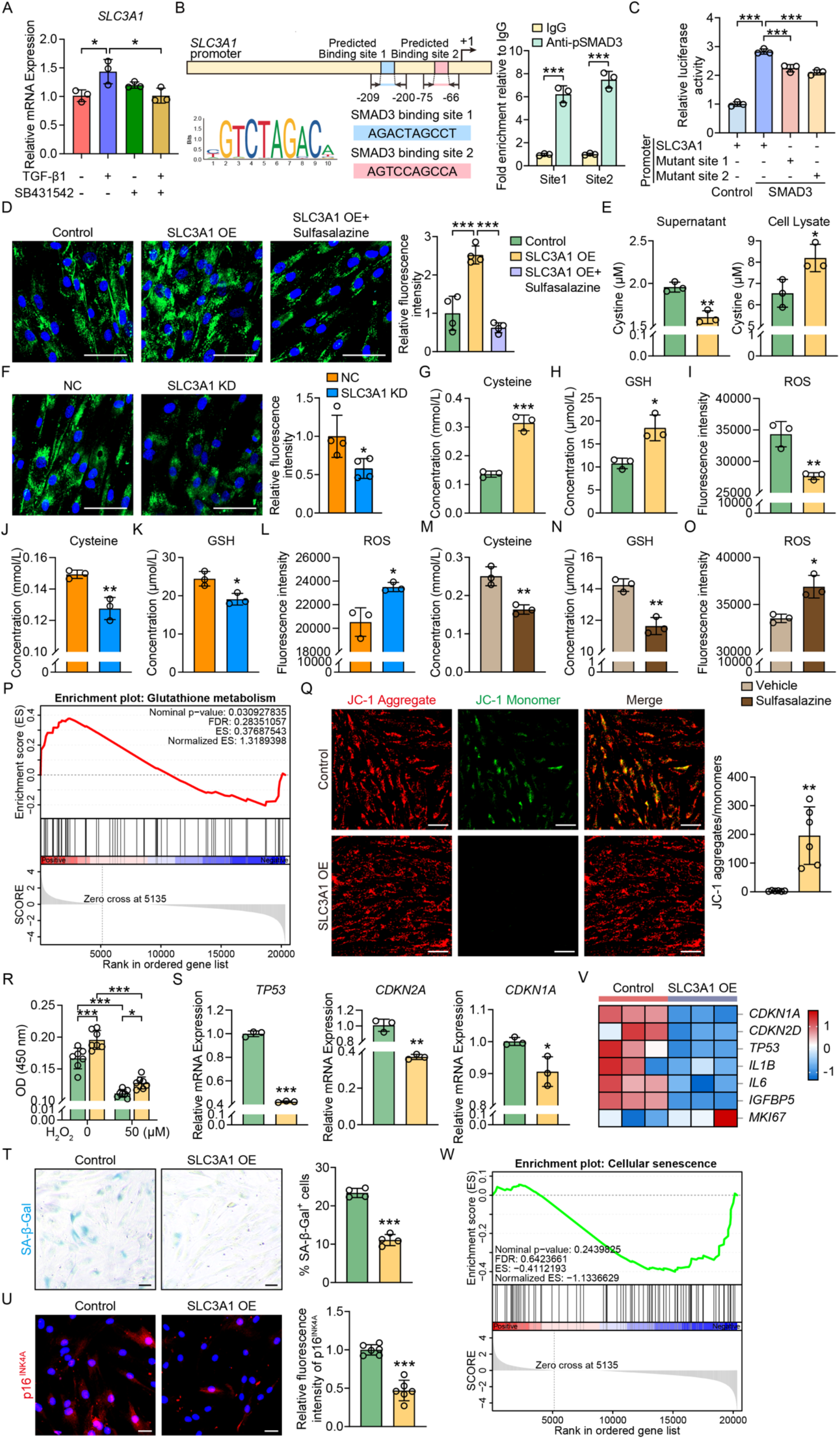
TGF-β-SMAD3 pathway upregulates SLC3A1 to activate cystine import and redox homeostasis to suppress stem cell senescence. **(A)** qPCR analysis showing that TGF-β1 stimulation (2 ng/mL) significantly upregulated *SLC3A1* mRNA levels in adipose-derived MSCs, while this induction was blocked by the TGF-β receptor inhibitor SB431542 (1 μM). **(B)** CUT&Tag analysis revealing SMAD3 enrichment at two predicted binding motifs in the *SLC3A1* promoter region. **(C)** Luciferase reporter assay showing that SMAD3 overexpression enhanced *SLC3A1* promoter activity, which was abrogated by site-directed mutagenesis of the SMAD3-binding motifs. **(D)** Representative images and quantification of FITC-labeled cystine uptake in control, SLC3A1-overexpressing, and Sulfasalazine-treated cells. **(E)** ELISA quantification showing increased intracellular and decreased extracellular cystine concentrations following SLC3A1 overexpression. **(F)** Representative images and quantification of FITC-cystine uptake in MSCs with SLC3A1 knockdown. **(G-I)** Biochemical measurements of intracellular cysteine **(G)**, glutathione (GSH, **H**), and reactive oxygen species (ROS, **I**) levels in control and SLC3A1-overexpressing cells. **(J-L)** Quantification of intracellular cysteine **(J)**, GSH **(K)**, and ROS **(L)** levels in negative control (NC) and SLC3A1-silenced cells. **(M-O)** Quantification of intracellular cysteine **(M)**, GSH **(N)**, and ROS **(O)** levels in vehicle and Sulfasalazine-treated cells. **(P)** GSEA analysis showing activation of glutathione metabolism in SLC3A1-overexpressing MSCs. **(Q)** JC-1 staining revealed increased mitochondrial membrane potential in SLC3A1-overexpressing MSCs, as indicated by higher red-to-green fluorescence ratio. **(R,S)** CCK-8 proliferation assay **(R)** and qPCR of senescence-associated genes *TP53*, *CDKN2A*, and *CDKN1A* **(S)** showing enhanced proliferative capacity and reduced senescence upon SLC3A1 overexpression. **(T)** SA-β-Gal staining showing decreased percentage of senescent cells in the SLC3A1-overexpressing group. **(U)** Immunofluorescence staining of p16 demonstrating reduced expression in SLC3A1-overexpressing cells. **(V,W)** RNA-seq heatmap **(V)** and GSEA analysis **(W)** confirming downregulation of senescence-related genes and the cellular senescence pathway in SLC3A1-overexpressing cells. Cell nuclei were counterstained with DAPI (blue). OE, overexpression; KD, knockdown. Scale bars: 50 μm. Data are presented as mean ± SD. \**P*<0.05; \*\**P*<0.01; \*\*\**P*<0.001.

**Figure 4.**
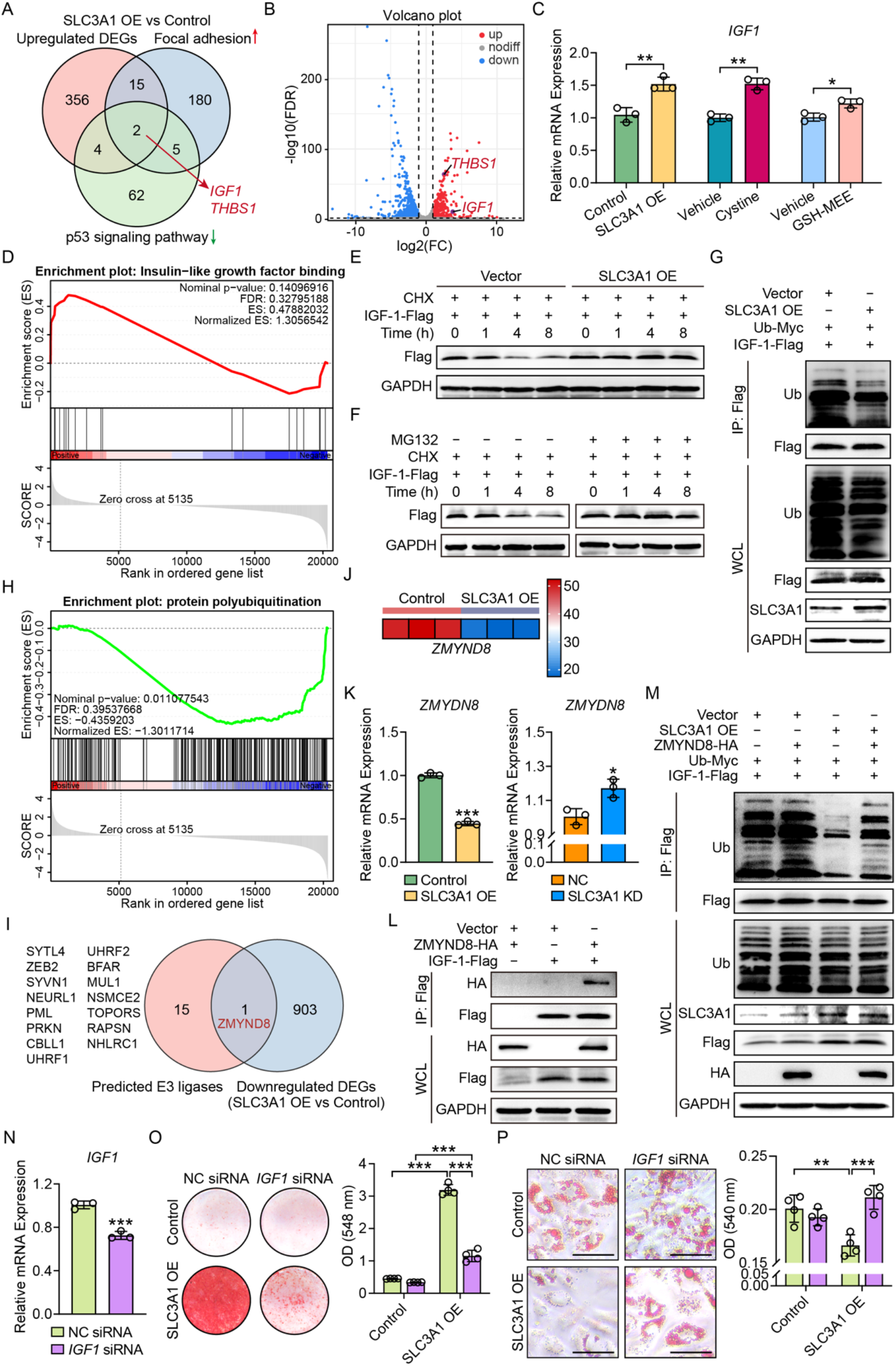
SLC3A1 enhances IGF-1 protein stability to drive osteogenic differentiation. **(A)** Venn diagram showing the gene overlap among three comparisons, upregulated DEGs from RNA-seq, and genes enriched in the Focal adhesion (activation) and p53 signaling pathway (inhibition) from KEGG analysis. Detailed gene lists were provided in **Supplementary Tables S4** and **S5**. **(B)** Volcano plot highlighting DEGs in control and SLC3A1-overexpressing MSCs (red: upregulated in SLC3A1 OE, blue: downregulated in SLC3A1 OE; FDR < 0.05, |log_2_FC| > 1). **(C,D)** qPCR and GSEA analysis confirmed the upregulation of *IGF1* and activation of insulin-like growth factor binding in SLC3A1-overexpressing cells. **(E)** SLC3A1 inhibited the degradation of IGF-1 protein. HEK293T cells were transfected with either empty vector or an SLC3A1-expressing plasmid, followed by treatment with cycloheximide (CHX) for the indicated times. Protein levels of IGF-1-Flag were detected by western blot. **(F)** The proteasome inhibitor inhibited the degradation of IGF-1 protein. HEK293T cells were treated with CHX alone or with proteasome inhibitor MG132 for the indicated times. **(G)** Representative western blots showing ubiquitination of IGF-1 in HEK293T cells transfected with empty vector or an SLC3A1-expressing plasmid. IGF-1 protein was immunoprecipitated (IP) with an anti-Flag antibody and detected with an anti-ubiquitin antibody. **(H)** GSEA analysis showing inhibition of protein polyubiquitination in SLC3A1 OE group. **(I)** Venn diagram showing the overlap between predicted E3 ligases for IGF-1 and downregulated DEGs from RNA-seq. **(J)** Heatmap revealed the expression levels of *ZMYND8* in control and SLC3A1 OE groups. **(K)** qPCR analysis of *ZMYND8* in control, SLC3A1-overexpressing, or SLC3A1-silenced MSCs. **(L)** Co-IP and western blot analysis confirmed the binding between ZMYND8 and IGF-1. **(M)** ZMYND8 promoted the ubiquitination of IGF-1 and reversed the stabilizing effect of SLC3A1. Representative western blots showing ubiquitination levels of IGF-1 under the indicated conditions. **(N)** qPCR analysis of *IGF1* knockdown efficiency. **(O)** Alizarin Red S staining and quantitative analysis of osteogenic differentiation following SLC3A1 overexpression and *IGF1* knockdown in four indicated groups. **(P)** Oil Red O staining and quantitative analysis of adipogenic differentiation following SLC3A1 overexpression and *IGF1* knockdown in four indicated groups. OE, overexpression; KD, knockdown; NC, negative control; Ub, ubiquitin. Scale bars: 50 μm. Data are presented as mean ± SD. \**P*<0.05; \*\**P*<0.01; \*\*\**P*<0.001.

We next examined whether SMAD3 directly regulates SLC3A1 transcription. *In silico* analysis identified two putative SMAD3-binding motifs within the *SLC3A1* promoter region. CUT&Tag analysis confirmed SMAD3 occupancy at both predicted binding sites (**Fig. 3B**). Moreover, luciferase reporter assays revealed that overexpression of SMAD3 enhanced *SLC3A1* promoter activity, whereas site-directed mutagenesis of the SMAD3-binding motifs significantly attenuated this activation (**Fig. 3C**). Altogether, these findings identify TGF-β/SMAD3 as a direct upstream regulator of SLC3A1 transcription and connect canonical osteogenic signaling to metabolic control of osteogenic fate.

### SLC3A1 enhances mitochondrial function and suppresses stem cell senescence

Given that SLC3A1 functions as a membrane transporter for cystine uptake^28,29^, we next investigated whether SLC3A1 regulates intracellular cystine metabolism and redox homeostasis. FITC-labeled cystine uptake assays demonstrated significantly increased intracellular cystine accumulation in SLC3A1-overexpressing mesenchymal cells, whereas cystine uptake was markedly reduced following *SLC3A1* knockdown or pharmacological inhibition with Sulfasalazine (**Fig. 3D,F**). Consistent with these findings, ELISA-based measurements showed reduced extracellular cystine levels accompanied by increased intracellular cystine concentrations in SLC3A1-overexpressing cells (**Fig. 3E**), confirming enhanced transporter activity. Since intracellular cystine is rapidly converted to cysteine, the precursor for glutathione (GSH) biosynthesis, we next examined the impact of SLC3A1 on cysteine-GSH metabolism. SLC3A1 overexpression significantly increased intracellular cysteine and GSH levels while reducing reactive oxygen species (ROS), as determined by biochemical assays and ROS-sensitive fluorescent probes. In contrast, *SLC3A1* knockdown or pharmacological blockade resulted in decreased cysteine and GSH abundance and elevated ROS accumulation (**Fig. 3G-O**). Consistent with these observations, transcriptomic analysis identified glutathione metabolism as one of the most significantly enriched pathways downstream of SLC3A1 activation (**Fig. 3P**). Collectively, these findings demonstrate that SLC3A1 promotes a redox-adaptive metabolic state through enhanced cystine utilization and glutathione production.

Given the established role of oxidative stress in mitochondrial dysfunction and cellular senescence^30–32^, we next evaluated whether SLC3A1-mediated redox adaptation influences cellular fitness. JC-1 staining revealed that SLC3A1 overexpression significantly enhanced mitochondrial membrane potential, indicating improved mitochondrial integrity and function (**Fig. 3Q**). Moreover, SLC3A1-overexpressing cells exhibited increased proliferative capacity as measured by CCK-8 assays and reduced expression of senescence-associated genes, including *TP53*, *CDKN2A*, and *CDKN1A* (**Fig. 3R,S**). These findings were further supported by decreased senescence-associated β-galactosidase (SA-β-Gal) positivity (**Fig. 3T**) and reduced p16 expression detected by immunofluorescence staining (**Fig. 3U**). At the transcriptomic level, senescence-associated pathways and gene signatures were broadly suppressed following SLC3A1 overexpression (**Fig. 3V,W**). In summary, these results identify SLC3A1 as a central regulator of cystine-dependent redox adaptation that preserves mitochondrial fitness and protects stem cells from senescence-associated functional decline.

### SLC3A1 stabilizes IGF-1 protein via ubiquitination blockade to promote osteogenesis

To identify molecular effectors linking SLC3A1-dependent metabolic adaptation to osteogenic fate regulation, we performed an integrative analysis by intersecting genes differentially expressed following SLC3A1 overexpression with pathways associated with cellular proliferation and senescence. This approach identified two candidate genes, *IGF1* and *THBS1* (**Fig. 4A**). Among them, *IGF1* showed robust upregulation in response to SLC3A1 overexpression (**Fig. 4B,C and Supplementary Fig. 5**). GSEA further demonstrated significant enrichment of insulin-like growth factor binding pathway in SLC3A1-overexpressing cells (**Fig. 4D**). Consistently, multiple components of the IGF-1 signaling network, including *IGF1* and its binding partner *IGFBP3*, were upregulated following SLC3A1A1 overexpression (**Fig. 4C and Supplementary Fig. 6**). Because signaling activity is frequently regulated through post-translational mechanisms, we next investigated whether SLC3A1 affects IGF-1 protein stability. Cycloheximide (CHX) chase assays demonstrated rapid degradation of IGF-1 protein under basal conditions, whereas SLC3A1 overexpression markedly prolonged IGF-1 protein stability (**Fig. 4E**). Treatment with the proteasome inhibitor MG132 prevented IGF-1 degradation, suggesting regulation through ubiquitin-dependent proteasomal turnover (**Fig. 4F**). Consistent with this model, co-immunoprecipitation (Co-IP) assays revealed substantially reduced ubiquitination of IGF-1 following SLC3A1 overexpression (**Fig. 4G**). Moreover, GSEA demonstrated suppression of protein polyubiquitination pathway in SLC3A1-overexpressing cells (**Fig. 4H**). Altogether, these findings suggest that SLC3A1 stabilizes IGF-1 by limiting ubiquitin-mediated degradation.

To identify the E3 ubiquitin ligase responsible for IGF-1 turnover, we performed computational prediction analysis and intersected the candidate ligases with genes downregulated following SLC3A1 overexpression. This analysis identified ZMYND8 as a putative regulator of IGF-1 stability (**Fig. 4I,J**). qPCR analysis confirmed reciprocal regulation between SLC3A1 and ZMYND8, with SLC3A1 overexpression suppressing ZMYND8 expression and SLC3A1 knockdown inducing ZMYND8 expression (**Fig. 4K**). Co-IP assays further demonstrated direct interaction between ZMYND8 and IGF-1 (**Fig. 4L and Supplementary Fig. 7**). Importantly, enforced ZMYND8 expression markedly increased IGF-1 ubiquitination (**Fig. 4M**), establishing ZMYND8 as a previously unrecognized E3 ubiquitin ligase that targets IGF-1 for degradation. These results identify a regulatory mechanism through which SLC3A1 preserves IGF-1 abundance by suppressing ZMYND8-dependent ubiquitination.

Finally, to determine whether IGF-1 mediates the osteogenic effects of SLC3A1, we silenced *IGF1* in SLC3A1-overexpressing mesenchymal cells (**Fig. 4N**). *IGF1* knockdown substantially attenuated SLC3A1-induced osteogenic differentiation, as evidenced by reduced mineralized matrix formation (**Fig. 4O**). In contrast, knockdown of *IGF1* restored adipogenic differentiation that had been suppressed by SLC3A1 activation (**Fig. 4P**). These findings establish IGF-1 as a critical downstream effector of SLC3A1 signaling. Collectively, these results identify a previously unrecognized SLC3A1-ZMYND8-IGF-1 regulatory axis in which SLC3A1 stabilizes IGF-1 protein by suppressing ZMYND8-mediated ubiquitination, thereby promoting osteogenic commitment.

### SLC3A1 activation enhances skeletal regeneration *in vivo*

To determine whether activation of SLC3A1 improves regenerative function *in vivo*, we generated MSCs stably overexpressing SLC3A1 (SLC3A1 OE) and verified successful transduction by fluorescence imaging and qPCR (**Supplementary Fig. 8**). These cells were combined with β-tricalcium phosphate (β-TCP) scaffolds and implanted subcutaneously into the dorsal region of nude mice to assess ectopic bone formation (**Fig. 5A**). After 8 weeks, micro-computed tomography (micro-CT) analysis revealed that SLC3A1 OE significantly enhanced bone regeneration, as evidenced by increased bone volume fraction (BV/TV), bone surface density (BS/TV), and trabecular number (Tb.N), accompanied by reduced trabecular separation (Tb.Sp) (**Fig. 5B,C**). Histological staining confirmed greater new bone deposition in the SLC3A1 OE group, with pronounced alkaline phosphatase (ALP) activity (**Fig. 5D,E**). GFP fluorescence confirmed the long-term local persistence of transplanted cells at the implantation site (**Fig. 5F**). Consistent with the cellular phenotypes observed *in vitro*, SLC3A1 overexpression reduced cell senescence and increased IGF-1 expression *in vivo* (**Fig. 5G,H**). Immunofluorescence staining further demonstrated elevated expression of both SLC3A1 and osteocalcin (OCN) in the SLC3A1 OE group, with clear co-localization between the two signals, indicating enhanced osteogenic activity within newly formed bone tissue (**Fig. 5I**). Therefore, SLC3A1-driven cellular reprogramming promotes a pro-regenerative osteogenic microenvironment *in vivo*.

**Figure 5.**
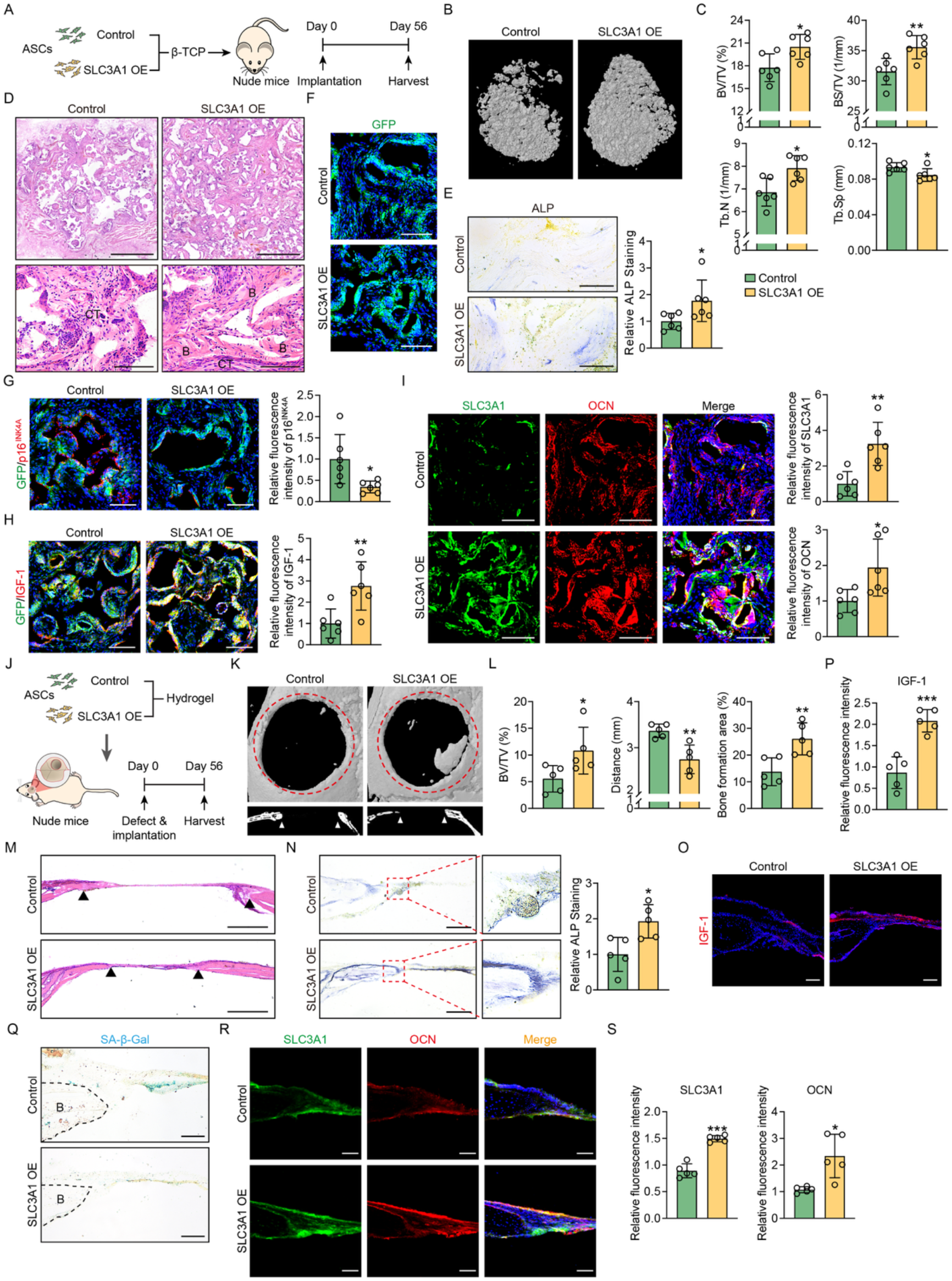
SLC3A1 overexpression enhances the osteogenic competence and regenerative function of MSCs *in vivo*. **(A)** Schematic illustration of the ectopic bone formation model. Adipose-derived MSCs with or without SLC3A1 overexpression were seeded onto β-TCP scaffolds and implanted subcutaneously into the dorsal region of nude mice. **(B)** Representative micro-CT images of explanted constructs at 8 weeks, demonstrating more robust bone formation in the SLC3A1 OE group. **(C)** Quantification of bone parameters by micro-CT analysis. SLC3A1 overexpression significantly increased bone volume fraction (BV/TV), bone surface density (BS/TV), and trabecular number (Tb.N), while reducing trabecular separation (Tb.Sp). **(D)** H&E staining of the implants. **(E)** Alkaline phosphatase (ALP) staining of explanted sections showing higher ALP activity in the SLC3A1 OE group. **(F)** The implants exhibited a robust presence of GFP-labeled cells, confirming the persistence of the exogenously delivered cells. **(G,H)** Representative immunofluorescence staining and quantification of p16 (senescence marker, **G**) and IGF-1 **(H)** in explanted sections. **(I)** Immunofluorescence co-staining indicating increased expression of SLC3A1 and osteocalcin (OCN) in the SLC3A1 OE group. **(J)** Schematic illustration of calvarial defect repair in nude mice. MSCs, either control or overexpressing SLC3A1, were encapsulated within a hydrogel scaffold and implanted into critical-sized defects in the parietal bone. After 8 weeks, skulls were harvested for the evaluation of bone formation. **(K)** Representative micro-CT 3D reconstructions of calvarial defects. Red circles mark the boundaries of the original defect sites. **(L)** Quantification of bone regeneration based on BV/TV, residual defect distance, and bone formation area. **(M)** H&E staining of the defect region showing tissue morphology. **(N)** Representative images of ALP staining and corresponding quantification of ALP activity. **(O,P)** Immunofluorescence staining and quantification of IGF-1 at the edge of the bone defect. **(Q)** SA-β-Gal staining revealed cellular senescence at the defect site. Black dashed lines indicate the defect margins. **(R)** Representative immunofluorescence images of SLC3A1 and OCN at the defect margins. **(S)** Quantitative analysis of SLC3A1 and OCN fluorescence intensity. Cell nuclei were counterstained with DAPI (blue). B, bone; CT, connective tissue; OE, overexpression. *n* = 5-6 per group. Scale bars: 1 mm (**D** (tile scan)), 500 μm (**M**), 200 μm (**E,N**), 100 μm (**D,F,G,H,I,O,Q,R**). Data are presented as mean ± SD. \**P*<0.05; \*\**P*<0.01; \*\*\**P*<0.001.

We next evaluated the regenerative efficacy of SLC3A1-overexpressing cells in a critical-sized calvarial defect model (**Fig. 5J**). At 8 weeks post-implantation, micro-CT analyses demonstrated markedly improved defect bridging in the SLC3A1 OE group, characterized by increased BV/TV and bone formation area, along with reduced residual defect distance (**Fig. 5K,L**). Histological examination confirmed enhanced bone regeneration and elevated ALP activity within the defect region (**Fig. 5M,N**). Consistent with findings from the ectopic bone formation model, SLC3A1 overexpression reduced cellular senescence, elevated IGF-1 levels, and promoted OCN expression *in vivo* (**Fig. 5O-S**). Collectively, these findings demonstrate that activation of SLC3A1 enhances osteogenic competence and promotes skeletal regeneration in both ectopic and orthotopic settings.

### Cystine supplementation phenocopies SLC3A1 activation and enhances skeletal regeneration

Given that SLC3A1 functions as a cystine transporter, we next asked whether exogenous cystine supplementation could recapitulate the cellular effects of SLC3A1 activation. Interestingly, cystine treatment itself significantly induced SLC3A1 expression (**Fig. 6A**), suggesting the existence of a positive feedback mechanism that reinforces cystine utilization. Consistent with enhanced transporter activity, cystine supplementation markedly increased intracellular cysteine and GSH levels while reducing ROS accumulation (**Fig. 6B**). Functionally, cystine-treated MSCs exhibited improved mitochondrial activity, enhanced proliferative capacity, and reduced cellular senescence, as evidenced by decreased expression of senescence-related genes and lower β-galactosidase and p16 activity (**Fig. 6C-G**). Moreover, cystine supplementation promoted osteogenic differentiation while suppressing adipogenesis (**Fig. 6H-K**). At the mechanistic level, cystine treatment effectively attenuated IGF-1 protein degradation, whereas knockdown of *IGF1* partially abolished the pro-osteogenic effects of cystine (**Fig. 6L,M**), further supporting IGF-1 as a critical downstream effector of the SLC3A1-cystine axis. Although SLC3A1 is capable of transporting other dibasic amino acids, including arginine, lysine, and ornithine^33,34^, supplementation with these substrates exerted only modest effects on osteogenic differentiation compared with cystine treatment (**Supplementary Fig. 9**). Among all amino acids tested, cystine exhibited the strongest pro-osteogenic activity, indicating that cystine acts as the dominant metabolic mediator of SLC3A1-dependent osteogenic programming. Overall, these findings demonstrate that cystine supplementation largely phenocopies the molecular and functional effects of SLC3A1 activation. Modulation of cystine availability represents a practical and effective strategy to metabolically reprogram stem cells toward an osteogenic state, providing a non-genetic approach to enhance regenerative function and therapeutic potential.

**Figure 6.**
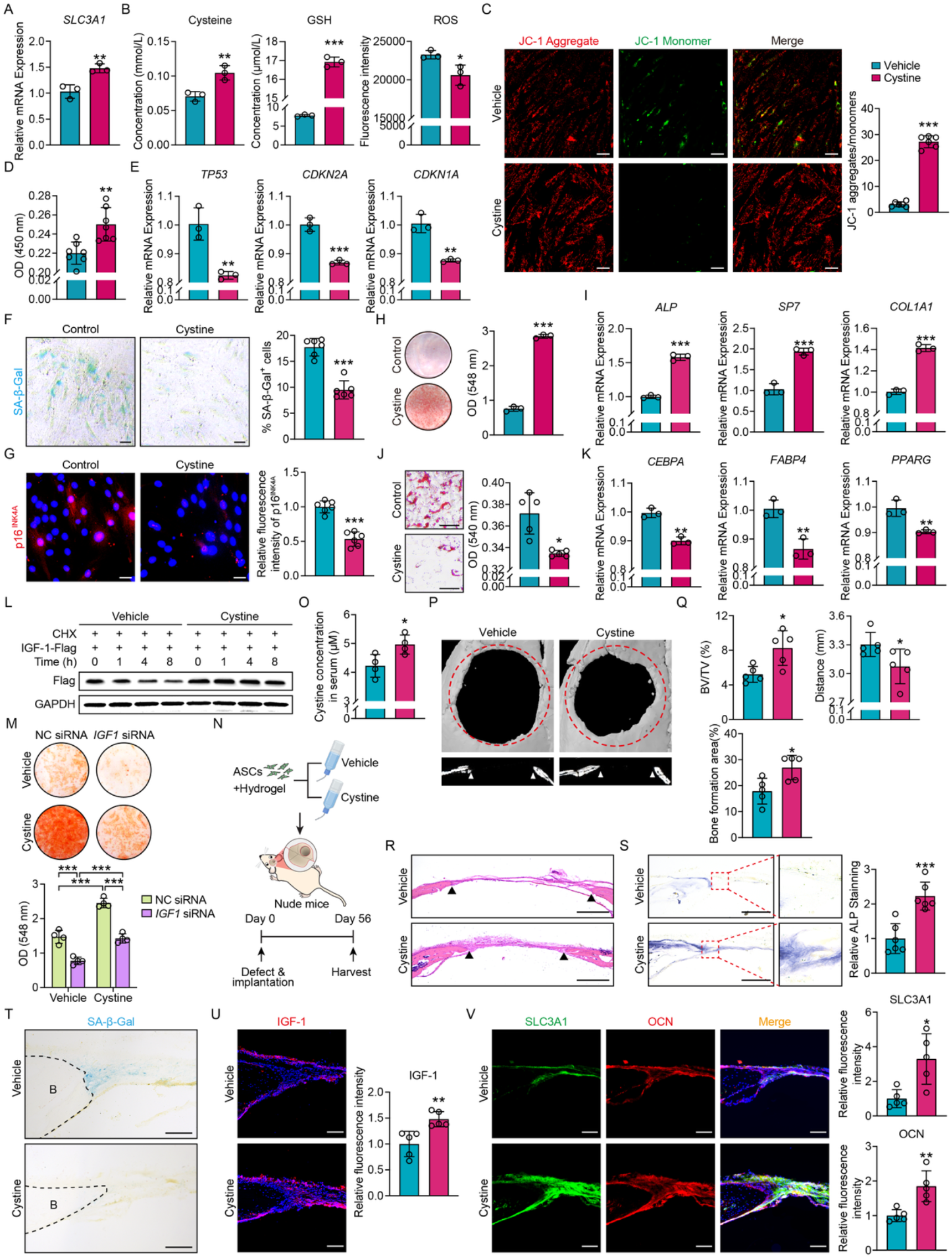
Cystine supplementation recapitulates the molecular and functional effects of SLC3A1 overexpression. **(A)** qPCR analysis of *SLC3A1* expression in adipose-derived MSCs following vehicle or cystine stimulation. **(B)** Intracellular levels of cysteine, GSH, and ROS were measured in cells treated with vehicle or cystine. **(C)** Cystine treatment enhanced mitochondrial membrane potential in MSCs, as shown by JC-1 staining. **(D)** CCK-8 assay showing increased proliferation. **(E)** qPCR showing downregulation of senescence markers (*TP53*, *CDKN2A*, *CDKN1A*) in cystine-treated cells. **(F)** SA-β-Gal staining revealed a reduced proportion of senescent cells in the cystine-treated group. **(G)** Immunofluorescence staining of p16 showed decreased expression following cystine treatment. **(H,I)** Alizarin Red S staining **(H)** and qPCR analysis **(I)** revealed enhanced osteogenic differentiation in MSCs following cystine treatment. **(J)** Oil Red O staining demonstrated reduced lipid droplet accumulation in MSCs treated with cystine. **(K)** qPCR analysis showed downregulation of adipocyte-associated genes (*CEBPA*, *FABP4*, *PPARG*) in cystine-stimulated group compared to vehicle control. **(L)** Cystine treatment reversed the degradation of IGF-1 protein. HEK293T cells were treated with either vehicle or cystine, followed by CHX treatment for the indicated time periods. Protein levels of IGF-1-Flag were detected by western blot. **(M)** Alizarin Red S staining and quantitative analysis of osteogenic differentiation following cystine treatment and *IGF1* knockdown in four indicated groups. **(N)** Schematic diagram of the calvarial defect model in nude mice. Hydrogel-embedded MSCs were implanted into critical-size defects in the parietal bone. Mice received vehicle or cystine through drinking water. Skulls were collected at 8 weeks for bone regeneration analysis. **(M)** Quantification of cystine concentration in mouse serum by ELISA. **(O)** ELISA analysis of serum cystine concentrations in peripheral blood collected from mice in the vehicle and cystine treatment groups. **(P)** Representative micro-CT 3D reconstructions of calvarial defects. Red circles indicate the boundaries of the original defect sites. **(Q)** Quantification of bone parameters derived from micro-CT, including BV/TV, residual defect distance, and bone formation area. **(R)** H&E-stained images of bone defect regions at 8 weeks post-implantation. **(S)** Representative images of ALP staining and quantification of ALP activity. **(T)** SA-β-Gal staining revealed cellular senescence at the defect site, with black dashed lines indicating the defect margins. **(U)** Immunofluorescence staining and quantitative analysis of IGF-1 expression at the edge of the bone defect. **(V)** Quantification of fluorescence intensity for SLC3A1 and OCN to assess their expression levels in the defect area. Cell nuclei were counterstained with DAPI (blue). B, bone. *n* = 5 per group. Scale bars: 500 μm (**R**), 200 μm (**S**), 100 μm (**T,U,V**), 50 μm (**C,F,G,J**). Data are presented as mean ± SD. \**P*<0.05; \*\**P*<0.01; \*\*\**P*<0.001.

Because cystine is already used clinically in other indications^35,36^, we next evaluated its regenerative efficacy *in vivo*. MSCs were implanted into critical-sized calvarial defects, and mice received cystine supplementation through drinking water (**Fig. 6N**). ELISA confirmed a significant increase in serum cystine levels in treated animals (**Fig. 6O**). Importantly, micro-CT and histological analyses revealed no evidence of renal or bladder stone formation (**Supplementary Fig. 10**), supporting the safety of this treatment regimen. Cystine supplementation markedly accelerated bone defect healing, as demonstrated by increased BV/TV and bone formation area, along with reduced defect width (**Fig. 6P,Q**). Histological analysis (H&E) corroborated these findings, revealing greater new bone formation and shortened defect distance in the cystine group (**Fig. 6R**). ALP staining further demonstrated elevated osteogenic activity (**Fig. 6S**), while SA-β-Gal staining and immunofluorescence analyses confirmed reduced cellular senescence, increased IGF-1 expression, and higher SLC3A1 and OCN expression at the repair site (**Fig. 6T-V**). Thus, these results provide strong evidence that cystine supplementation is sufficient to reproduce key biological effects of SLC3A1 activation and promotes skeletal regeneration through enhancement of osteogenic competence.

### Cystine supplementation promotes endogenous bone formation and attenuates osteoporotic bone loss

To determine whether the pro-osteogenic effects of the SLC3A1-cystine axis are conserved across mesenchymal stem/stromal cell populations, we overexpressed SLC3A1 in bone marrow-derived MSCs and evaluated lineage differentiation. Similar to the findings observed in adipose-derived cells, SLC3A1 overexpression enhanced osteogenic differentiation, increased expression of osteogenic markers, and suppressed adipogenic differentiation and lipid accumulation (**Fig. 7A-D**). These results demonstrate the functional generalizability of SLC3A1 in promoting osteogenic differentiation across distinct mesenchymal stem/stromal cell types, underscoring its potential as a universal target for skeletal regeneration and bone disease therapy.

**Figure 7.**
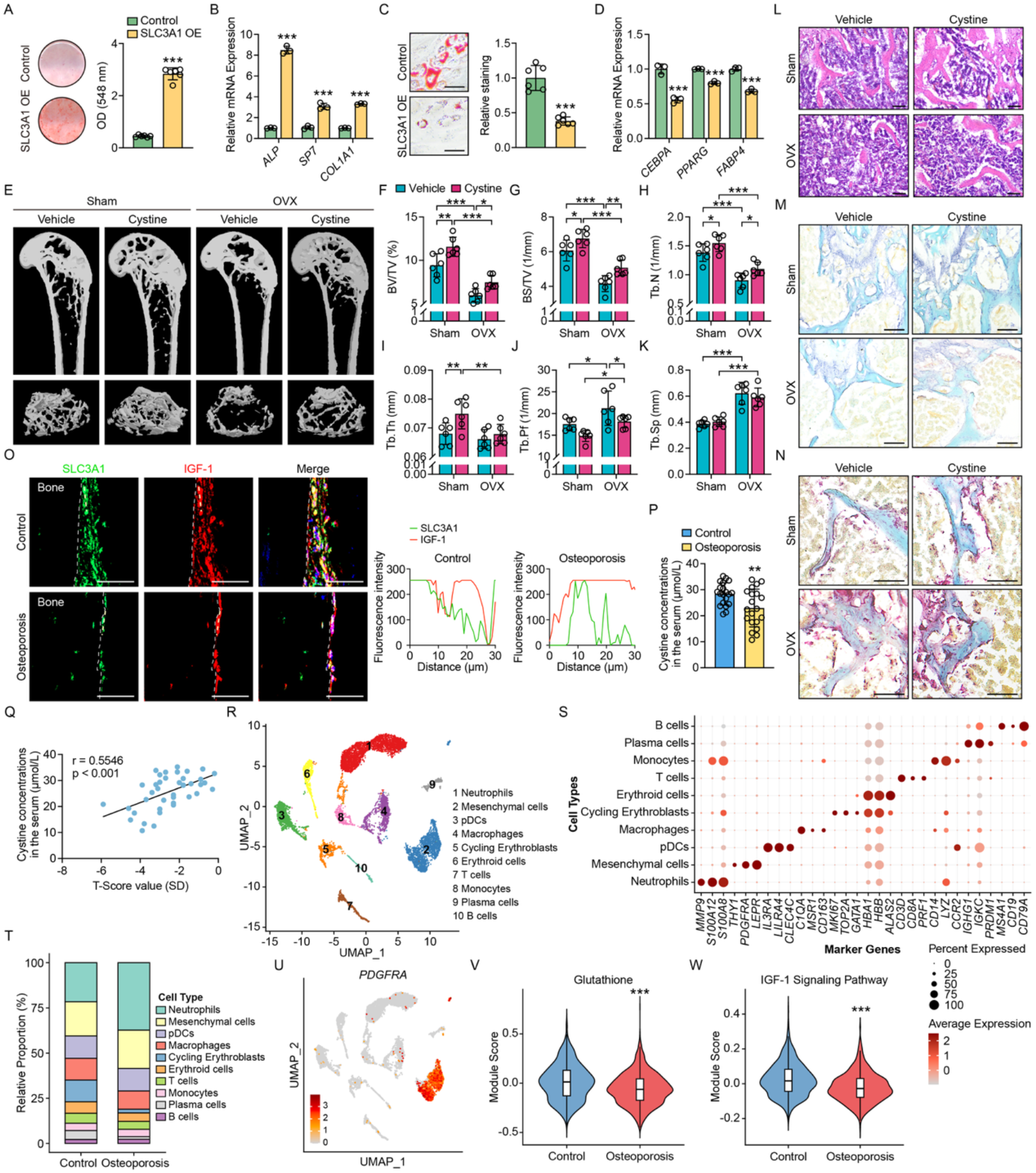
SLC3A1 overexpression enhances bone marrow MSC osteogenesis and cystine supplementation prevents bone loss in OVX mice. **(A)** Alizarin Red S staining showing increased mineralized nodule formation in SLC3A1-overexpressing bone marrow MSCs following osteogenic induction. **(B)** qPCR analysis of osteogenic gene expression (*ALP*, *SP7*, *COL1A1*) in control and SLC3A1-overexpressing cells. **(C)** Oil Red O staining showing reduced lipid droplet accumulation in SLC3A1 OE group under adipogenic conditions. **(D)** qPCR analysis of adipogenic markers (*CEBPA*, *PPARG*, *FABP4*). **(E)** Representative micro-CT images of distal femurs from sham and OVX mice treated with vehicle or cystine for 8 weeks. *n* = 6 per group. **(F-K)** Quantification of micro-CT-derived trabecular bone parameters, including **(F)** BV/TV, **(G)** BS/TV, **(H)** Tb.N, **(I)** Tb.Th, **(J)** Tb.Pf, and **(K)** Tb.Sp. **(L)** Representative H&E staining of femoral trabecular bone sections. **(M)** ALP staining indicating increased osteoblast activity upon cystine supplementation. **(N)** TRAP staining showing no significant difference in osteoclast numbers among groups. **(O)** Immunofluorescence staining of SLC3A1 and IGF-1 on bone surfaces in control and osteoporotic human samples, with co-localization analysis. White dashed lines indicate the bone surface boundary. **(P)** ELISA analysis of serum cystine concentrations in peripheral blood collected from control subjects and osteoporotic patients. *n* = 20 per group. **(Q)** Correlation analysis showing a positive association between serum cystine levels and T-score values, an indicator of bone mineral density. **(R)** Single-cell RNA-sequencing analysis of bone marrow mononuclear cells isolated from control and osteoporotic patients. UMAP visualization showing cellular clustering. **(S)** Dot plot showing cell type-specific marker gene expression across identified cell populations. **(T)** Relative proportions of different cell populations in control and osteoporotic samples. **(U)** UMAP visualization of *PDGFRA* expression identifying mesenchymal cell populations, which were subsequently subjected to pathway enrichment analysis. **(V,W)** AddModuleScore analysis showing reduced glutathione pathway activity **(V)** and suppressed IGF-1 signaling **(W)** in mesenchymal cells from osteoporotic samples. OE, overexpression. Scale bars: 50 μm (**C**), 100 μm (**L,M,N**), 50 μm (**O**). Data are presented as mean ± SD. \**P*<0.05; \*\**P*<0.01; \*\*\**P*<0.001.

We next evaluated whether dietary cystine supplementation could enhance endogenous bone regeneration *in vivo* using a murine tibial defect model. Micro-CT and histological analyses demonstrated accelerated bone repair in cystine-treated animals, characterized by increased BV/TV, BS/TV, and Tb.N, reduced Tb.Sp, and enhanced osteogenic activity within the defect region (**Supplementary Fig. 11A-G**). Immunofluorescence staining further revealed that cystine treatment elevated SLC3A1 and OCN expression and markedly increased their colocalization (**Supplementary Fig. 11H,I**), indicating that cystine effectively promotes endogenous stem cell osteogenic differentiation and bone regeneration *in vivo*.

To further assess the therapeutic relevance of cystine supplementation, we examined its effects in ovariectomy-induced osteoporosis. Micro-CT analysis showed that cystine not only increased trabecular bone mass in baseline mice, but also markedly attenuated ovariectomized (OVX)-induced bone loss, as evidenced by higher BV/TV, BS/TV, Tb.N, and trabecular thickness (Tb.Th), along with reduced trabecular pattern factor (Tb.Pf) and Tb.Sp (**Fig. 7E-K**). H&E staining revealed preservation of trabecular structure and bone mass in cystine-treated OVX mice (**Fig. 7L**). Moreover, ALP staining indicated enhanced osteogenic activity, whereas TRAP staining showed no significant change in osteoclast abundance (**Fig. 7M,N**), suggesting that cystine primarily acts by stimulating osteogenic lineage commitment of skeletal stem/progenitor cells rather than by inhibiting osteoclastogenesis.

Finally, we asked whether dysregulation of the SLC3A1-cystine pathway also occurs in human osteoporosis. Analysis of osteoporotic patient specimens revealed markedly reduced expression of both SLC3A1 and IGF-1, accompanied by diminished co-localization signals compared with non-osteoporotic controls (**Fig. 7O**). Consistently, serum cystine levels were significantly lower in osteoporotic patients and positively correlated with bone mineral density (**Fig. 7P,Q**), suggesting a close association between systemic cystine availability and skeletal health. To further assess the clinical relevance of this pathway, we analyzed publicly available single-cell transcriptomic datasets from human osteoporotic bone tissues, focusing on mesenchymal stromal populations (**Fig. 7R-U**). Mesenchymal cells from osteoporotic samples exhibited coordinated suppression of glutathione metabolic programs together with reduced IGF-1 signaling activity (**Fig. 7V,W**), indicating disruption of the SLC3A1-cystine-IGF-1 regulatory axis in human osteoporosis. These alterations closely mirrored the metabolic and transcriptional phenotypes observed in our *in vitro* and murine studies, supporting the clinical relevance of this metabolic program in human skeletal aging and degeneration. Collectively, these findings support dietary cystine supplementation as a simple, safe, and clinically translatable strategy to enhance endogenous bone formation and protect against osteoporosis.

## Discussion

In this study, we identified a previously unrecognized metabolic mechanism that governs osteogenic fate determination in mesenchymal stem/stromal cells. Through integrated transcriptomic, mechanistic, and functional analyses, we demonstrated that activation of the SLC3A1-cystine axis established a redox-adaptive metabolic state that preserved mitochondrial fitness, suppressed cellular senescence, and stabilized the pro-osteogenic regulator IGF-1 through inhibition of ZMYND8-mediated ubiquitination. These coordinated effects promoted osteogenic competence and enhanced skeletal regeneration across multiple experimental settings. Furthermore, dietary supplementation with cystine largely phenocopied the effects of SLC3A1 activation and improved bone regeneration in both injury and osteoporosis models. Therefore, these findings identify the SLC3A1-cystine axis as a central regulator of osteogenic fate and reveal a metabolically driven strategy for enhancing skeletal regeneration (**Fig. 8**).

**Figure 8.**
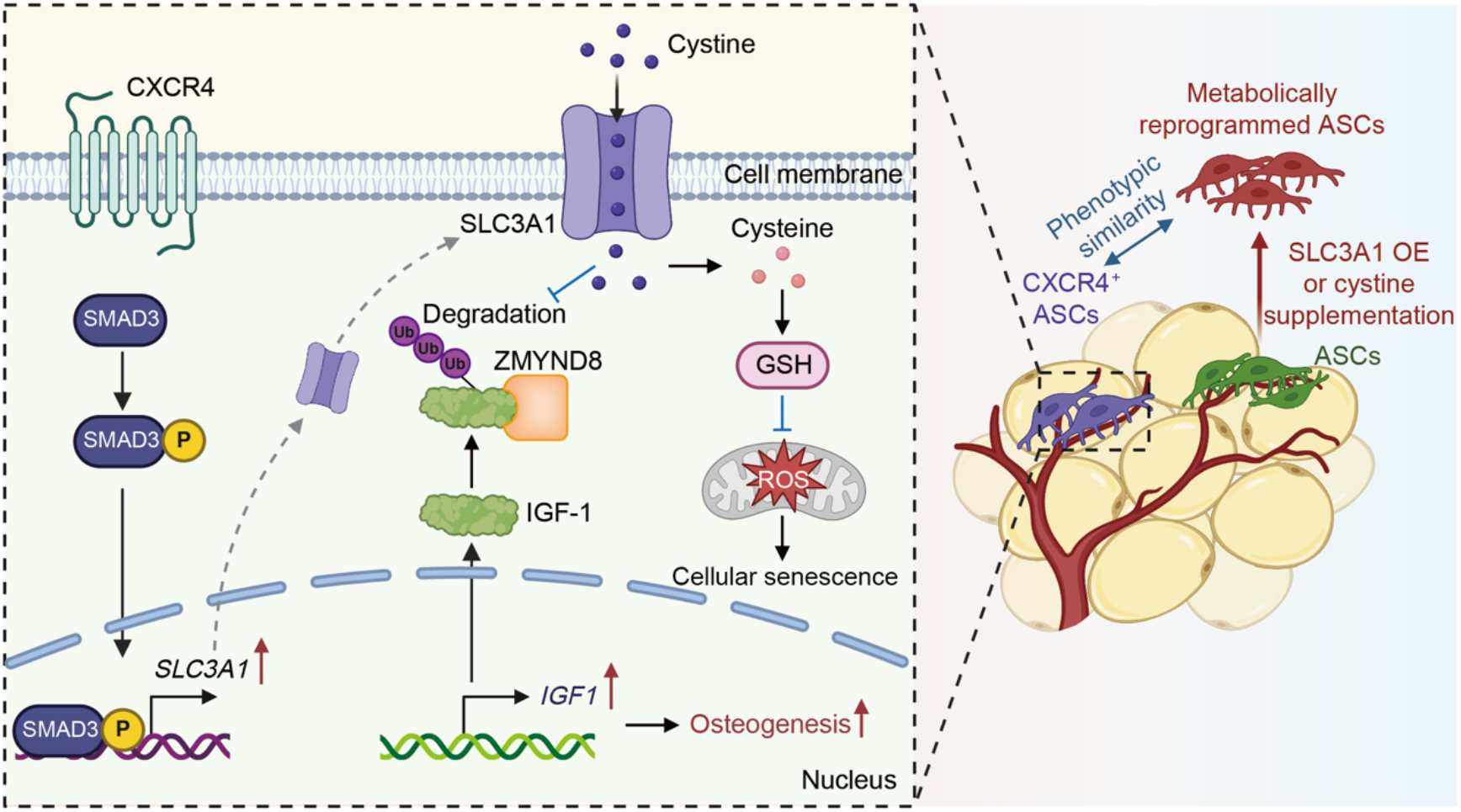
Schematic illustration of the SLC3A1-cystine axis in regulating osteogenic fate determination of MSCs. CXCR4^+^ mesenchymal cells exhibit elevated TGF-β/SMAD3 signaling, which directly activates *SLC3A1* transcription. Enhanced SLC3A1-mediated cystine uptake increases intracellular glutathione (GSH) levels, suppresses reactive oxygen species (ROS) accumulation, and establishes a redox-adaptive metabolic state that preserves mitochondrial fitness and limits cellular senescence. In parallel, SLC3A1 stabilizes the osteogenic regulator IGF-1 through suppression of ZMYND8-mediated ubiquitination and proteasomal degradation, thereby sustaining osteogenic commitment. Functionally, SLC3A1 overexpression (OE) or cystine supplementation metabolically reprograms bulk mesenchymal cells toward a CXCR4^+^-like osteogenic phenotype, resulting in enhanced osteogenic competence and skeletal regeneration. Red arrows indicate activation or enhancement, whereas blue lines indicate inhibition.

Our findings also provide new insights into the biological significance of the CXCR4^+^ mesenchymal cell population that we previously identified as possessing superior osteogenic potential^26,37^. Although CXCR4 has long been recognized as a surface marker for multiple stem and progenitor populations^38^, its mechanistic role in osteogenic regulation remains incompletely defined. Previous studies suggested that activation of the SDF-1/CXCR4 axis promotes osteoblast differentiation, whereas pharmacological inhibition with AMD3100 impairs bone formation, likely by blocking SDF-1 binding within the bone microenvironment. In contrast, recent evidence from WHIM syndrome, a rare immunodeficiency caused by gain-of-function CXCR4 mutations, reveals that 25% of patients exhibit low bone mass, and WHIM mutant mice develop osteopenia characterized by decreased osteoprogenitor frequency^39^. Constitutive CXCR4 activation disrupts bone-marrow stromal cell cycling and osteogenic differentiation. Long-term AMD3100 administration in this model restores osteogenesis and reverses bone loss, underscoring that precise tuning rather than constitutive activation of CXCR4 is critical for skeletal homeostasis. Interestingly, direct neutralization of CXCR4 in our study failed to impair the osteogenic potential of CXCR4^+^ cells. This finding diverges from earlier reports attributing a pro-osteogenic role to SDF-1/CXCR4 signaling^40,41^, but may reflect the absence of sustained SDF-1 stimulation *in vitro*. Rather than functioning as an active signaling effector, CXCR4 appears to serve predominantly as a phenotypic marker that identifies a mesenchymal cell population already primed toward osteogenesis. Consistent with this interpretation, CXCR4^+^ cells exhibited elevated TGF-β/SMAD3 activity and marked enrichment of SLC3A1 expression. Collectively, these observations support a model in which CXCR4 identifies osteogenic competence, whereas the underlying osteogenic state is governed by intracellular signaling and metabolic programs centered on the SLC3A1-cystine pathway.

Current approaches to enhance skeletal regeneration have largely focused on identifying osteogenic cell populations or manipulating lineage-specific transcriptional regulators. However, these strategies are frequently limited by cellular heterogeneity, loss of function during ex vivo expansion, and poor scalability. Our findings suggest an alternative framework in which osteogenic competence can be enhanced through metabolic adaptation rather than cell selection alone. Activation of the SLC3A1-cystine pathway increases intracellular glutathione availability, limits oxidative stress, preserves mitochondrial function, and stabilizes IGF-1 protein abundance, thereby establishing a cellular state that favors osteogenic commitment. This concept extends beyond conventional gene-centric models of lineage regulation and highlights metabolic state as an active determinant of regenerative function. Rather than merely supporting cellular bioenergetics^16^, amino acid metabolism can directly shape lineage commitment through coordinated regulation of redox homeostasis, protein stability, and cellular senescence.

An additional conceptual advance of this study is the identification of ZMYND8 as a previously unrecognized E3 ubiquitin ligase for IGF-1. Although IGF-1 signaling has been implicated in skeletal development and osteogenic differentiation^42–45^, mechanisms controlling IGF-1 protein stability have remained largely unknown. Our data demonstrated that ZMYND8 promoted IGF-1 ubiquitination and degradation, whereas activation of the SLC3A1 pathway preserved IGF-1 abundance by suppressing ZMYND8-dependent turnover. These findings establish a previously unrecognized SLC3A1-ZMYND8-IGF-1 regulatory axis linking metabolic status to osteogenic signaling and provide a mechanistic explanation for how metabolic adaptation can be translated into durable changes in cell fate.

The observation that dietary cystine supplementation largely phenocopied SLC3A1 activation further highlights the translational potential of this pathway. Unlike genetic manipulation, metabolic intervention provides a comparatively accessible strategy to enhance osteogenic competence and skeletal regeneration. Because cystine is already used clinically in other indications, the safety profile and translational feasibility of this approach may be advantageous. Moreover, the conserved effects observed in both adipose-derived and bone marrow-derived MSCs suggest that modulation of cystine metabolism may represent a broadly applicable strategy for regenerative and age-associated skeletal disorders.

An intriguing observation from this study is that cystine consistently exerted stronger biological effects than cysteine, despite both metabolites promoting osteogenic differentiation (**Supplementary Fig. 12 A-D**). Interestingly, cystine supplementation stabilized IGF-1 protein, whereas cysteine treatment failed to do so (**Supplementary Fig. 12E**). Because cystine is the preferred extracellular substrate of SLC3A1-mediated transport, these findings suggest that transporter-dependent cystine utilization may exert biological functions beyond simply supplying intracellular cysteine for glutathione synthesis. Such effects may involve distinct metabolic fluxes or signaling events that ultimately converge on regulation of protein stability and osteogenic fate determination.

Several limitations of this study should be acknowledged. First, although cystine supplementation demonstrated robust osteogenic benefits in our models, the long-term safety profile, optimal dosage, and potential off-target effects of chronic administration remain to be fully evaluated. Second, we did not employ MSC-specific SLC3A1 knockout mouse models, which would provide more definitive genetic evidence for the role of SLC3A1 in osteogenic regulation *in vivo*. Third, our findings were obtained primarily in rodent models. Validation in large animal models with bone defects, as well as clinical investigations such as monitoring bone mass in patients receiving cystine for other indications, will be essential to confirm translational potential. Future studies addressing these points will help to further clarify the mechanisms, refine the therapeutic strategy, and accelerate clinical application of SLC3A1/cystine-based interventions for bone regeneration and skeletal diseases.

In summary, this study identifies the SLC3A1-cystine axis as a metabolic gatekeeper of osteogenic fate. By coordinating redox adaptation, mitochondrial fitness, cellular senescence, and IGF-1 stability, this pathway establishes a metabolic framework that supports osteogenic competence and skeletal regeneration. These findings provide mechanistic insight into how metabolic state governs lineage commitment and suggest that targeting amino acid metabolism may offer a clinically accessible strategy for enhancing skeletal repair and combating osteoporosis.

## Materials and methods

### Mice

All animal procedures were conducted in compliance with the approved protocol of the Institutional Animal Care and Use Committee at Southern Medical University. Male nude mice and C57BL/6J mice were purchased from Zhuhai Bestest Biotechnology Co., Ltd. (Zhuhai, China).

### Isolation and culture of human mesenchymal stem/stromal cells (MSCs)

Human adipose tissue and femoral bone samples was obtained from adult donors undergoing elective surgery at Nanfang Hospital, Southern Medical University, with institutional ethics committee approval. Detailed donor information is provided in **Supplementary Table S6**.

For isolation of adipose-derived MSCs, adipose samples were minced and digested in Hank’s Balanced Salt Solution (HBSS) containing 1 mg/mL collagenase type II (Biosharp, Hefei, Anhui, China) and 0.5% bovine serum albumin (BSA; Biosharp) at 37°C for 1 hour. The resulting cell suspension was centrifuged to remove blood and lipid layers, treated with red blood cell lysis buffer (Solarbio, Beijing, China) for 5 minutes at room temperature (RT), filtered through a 70 μm cell strainer, and centrifuged to obtain the stromal vascular fraction. For isolation of bone marrow-derived MSCs, the bone marrow cavity of femoral specimens was repeatedly flushed with phosphate-buffered saline (PBS) using a sterile syringe. The flushed suspension was passed through a 70 μm cell strainer and centrifuged to collect nucleated cells. Cells isolated from both sources were cultured in α-MEM (Gibco, Grand Island, NY) supplemented with 10% fetal bovine serum (FBS; Gibco) and 1% penicillin-streptomycin (Gibco) at 37°C with 5% CO_2_. Culture medium was refreshed every 3 days, and cells were passaged at a 1:2 ratio upon reaching 80-90% confluency.

### Flow cytometry and cell sorting

Single-cell suspensions were centrifuged and resuspended in HBSS containing 0.5% BSA. Cells were incubated with fluorophore-conjugated antibodies for 20 minutes on ice in the dark. After staining, cells were washed twice with HBSS + 0.5% BSA by centrifugation and resuspended in the same buffer for analysis or sorting. Fluorescence-activated cell sorting (FACS) was performed using a MoFlo XDP cell sorter (Beckman Coulter, Indianapolis, IN), and analytical flow cytometry was conducted on a BD LSRFortessa cytometer (BD Biosciences, San Jose, CA). Data were analyzed using FlowJo software (TreeStar, Ashland, OR). Detailed antibody information is provided in **Supplementary Table S1**.

### Osteogenic differentiation

MSCs were seeded into 12- or 48-well plates at a density of 2 × 10^5^ cells/mL. After 24 hours, the growth medium was replaced with osteogenic differentiation medium (ODM) composed of α-MEM supplemented with 10% FBS, 1% penicillin-streptomycin, 10 mM β-glycerophosphate (Sigma-Aldrich, St. Louis, MO), 50 μM ascorbic acid (Sigma-Aldrich), and 100 nM dexamethasone (Sigma-Aldrich). The medium was refreshed every 3 days. After 6-14 days of induction, Alizarin Red S (AR) staining was performed to assess mineralization. Cells were first washed with PBS, fixed in 4% paraformaldehyde (PFA), and then stained with 2% AR solution for 5-10 minutes at RT. Excess dye was removed by washing with PBS. For quantification, the calcium-bound dye was eluted with 0.1 N NaOH, and absorbance was measured at 548 nm using a SpectraMax i3x Multi-Mode Microplate Reader (Molecular Devices, San Jose, CA).

### Adipogenic differentiation

MSCs were seeded into 12- or 48-well plates at a density of 2 × 10^5^ cells/mL and allowed to adhere overnight. The culture medium was replaced with adipogenic differentiation medium (ADM) composed of α-MEM, 10% FBS, 1% penicillin-streptomycin with 1 μM dexamethasone, 0.5 mM IBMX, 10 μg/mL insulin, and 200 μM indomethacin. The medium was refreshed every 3 days. After 10-14 days of induction, Oil Red O staining was performed to visualize intracellular lipid accumulation. Cells were washed with PBS, fixed in 4% PFA, and incubated with working Oil Red O solution (prepared by diluting a stock solution of 5 mg/mL Oil Red O in isopropanol with ddH_2_O at a 3:2 ratio) at 37°C for 30 minutes. After staining, cells were washed with PBS and observed under a microscope. Lipid-bound dye was then eluted with isopropanol, and absorbance was measured at 540 nm using a microplate reader.

### Quantitative polymerase chain reaction (qPCR)

Total RNA was extracted from cultured cells using TRIzol reagent (Vazyme, Nanjing, China), and complementary DNA (cDNA) was synthesized using HiScript III All-in-one RT SuperMix Perfect for qPCR (Vazyme) according to the manufacturer’s instructions. qPCR was performed using Taq Pro Universal SYBR qPCR Master Mix (Vazyme) on a QuantStudio 5 Real-Time PCR System (Applied Biosystems, Waltham, MA). Relative gene expression levels were calculated using the 2^−ΔΔCt^ method and normalized to GAPDH as the internal control. Primer sequences are listed in **Supplementary Table S2**.

### Co-immunoprecipitation (Co-IP)

HEK293T cells were cultured in complete DMEM medium and transiently transfected with plasmids encoding ZMYND8-HA, IGF-1-Flag, ubiquitin-Myc, and SLC3A1, as indicated for each experimental group. Three days after transfection, cells were harvested and lysed in ice-cold IP lysis buffer (Vazyme) supplemented with protease inhibitor cocktail (MedChemExpress, Monmouth Junction, NJ). Cell lysates were clarified by centrifugation, and a portion of the supernatant was collected as the input control. The remaining lysates were incubated with anti-Flag antibody at 4°C overnight, followed by incubation with protein A/G magnetic beads (MedChemExpress). Normal IgG was used as a negative control. After extensive washing with lysis buffer, the immunoprecipitated proteins were eluted by boiling in SDS loading buffer and subjected to SDS-PAGE and immunoblotting. The precipitated complexes were analyzed using antibodies against HA, Flag, and ubiquitin to detect ZMYND8-HA, IGF-1-Flag, and ubiquitinated proteins, respectively.

### Cycloheximide (CHX) chase assay

HEK293T cells were transiently transfected with IGF-1-Flag together with SLC3A1 or IGF-1-Flag alone, followed by treatment with or without cystine (1 mM) or N-acetylcysteine (NAC, 1 mM) as indicated. Three days after transfection, cells were treated with CHX (25 μg/mL) to block new protein synthesis and harvested at 0, 1, 4, and 8 h after CHX treatment. For proteasome inhibition experiments, cells were pretreated with MG132 (10 μM) for 8 h before CHX treatment and maintained in medium containing both CHX and MG132 during the chase period. Total cell lysates were collected at each time point and subjected to SDS-PAGE and immunoblotting. The protein level of IGF-1-Flag was detected using an anti-Flag antibody.

### Western blot

Cells were lysed in RIPA buffer supplemented with 1 mM PMSF (Beyotime Biotech Inc., Shanghai, China), and total protein concentration was determined using an enhanced BCA protein assay kit (Beyotime Biotech Inc.). The lysates were combined with 5× SDS-PAGE loading buffer and denatured by boiling at 100 °C for 10 minutes. Equal amounts of protein (20 μg per lane) were then resolved on 10% SDS-polyacrylamide gels. Proteins were transferred to polyvinylidene difluoride (PVDF) membranes (Millipore, Darmstadt, Germany), blocked with 5% BSA for 1 h at RT, and incubated with primary antibodies overnight at 4°C. After washing with TBST, membranes were incubated with horseradish peroxidase (HRP)-conjugated secondary antibodies for 1.5 h at RT. Signals were visualized using an enhanced chemiluminescence detection system (Vazyme), and densitometric analysis was performed using ImageJ software (National Institutes of Health, Bethesda, MD). Detailed antibody information is provided in **Supplementary Table S1**.

### CUT&Tag

CUT&Tag was performed according to the manufacturer’s protocol. In brief, 1 × 10^5^ MSCs were resuspended in wash buffer and incubated with 10 μL of pre-treated ConA Beads Pro (Vazyme) at RT for 10 minutes. After discarding the supernatant, bead-bound cells were resuspended in 50 μL of antibody buffer containing p-SMAD3 primary antibody (1:200; Cell Signaling Technology, Danvers, MA) and incubated overnight at 4 °C. The following day, goat anti-rabbit IgG secondary antibody and pA/G-Tnp Pro were sequentially added and incubated at RT for 1 hour each. Next, 10% SDS and 1 pg of DNA spike-in were added to the samples and incubated at 55 °C for 10 minutes. The supernatant was then collected and incubated with DNA Extract Beads Pro for 20 minutes. Beads were washed twice with B&W buffer and resuspended in ddH_2_O. To release the bound DNA, 5 μL of stop buffer was added to the bead suspension and heated at 95 °C for 5 minutes. The resulting DNA was analyzed by CUT&Tag-qPCR using Taq Pro Universal SYBR qPCR Master Mix (Vazyme). Relative enrichment was normalized to the DNA spike-in control. Primer sequences are listed in **Supplementary Table S2**.

### RNA Sequencing

Total RNA was extracted using TRIzol reagent (Vazyme) following the manufacturer’s instructions. Eukaryotic mRNA was enriched using Oligo(dT) beads, fragmented into short fragments with fragmentation buffer, and subsequently reverse-transcribed into cDNA using the NEBNext Ultra RNA Library Prep Kit for Illumina (New England Biolabs, Ipswich, MA). The resulting cDNA libraries were sequenced on the Illumina NovaSeq 6000 platform by Gene Denovo Biotechnology Co., Ltd. (Guangzhou, China). Principal component analysis (PCA) was conducted using the gmodels R package. Differential gene expression between groups was analyzed using DESeq2. Functional enrichment analyses, including Gene Ontology (GO), Kyoto Encyclopedia of Genes and Genomes (KEGG), and Gene Set Enrichment Analysis (GSEA), were performed to investigate the biological functions and signaling pathways associated with differentially expressed genes.

Single-cell RNA-sequencing data were analyzed using Seurat (v4.0) in R. Cells with low-quality features were excluded prior to normalization and downstream analysis. Highly variable genes were identified and used for dimensionality reduction and unsupervised clustering. Cellular heterogeneity and cluster distributions were visualized using Uniform Manifold Approximation and Projection (UMAP). Pathway activity and gene module enrichment scores were calculated using the AddModuleScore function in Seurat with curated gene signatures.

### Transfection

The pECMV-3×FLAG, pECMV-3×FLAG-SMAD3, pLV3-CMV-MCS-3×FLAG-CopGFP-Puro and pLV3-CMV-SLC3A1-3×FLAG-CopGFP-Puro plasmids were purchased from Miaoling Biotechnology Co., Ltd. (Wuhan, China). Plasmids or siRNAs were transfected into MSCs or HEK293T cells using Lipofectamine 2000 (ThermoFisher Scientific, Waltham, MA) in accordance with the manufacturer’s instructions. Four hours post-transfection, the transfection reagent was removed and replaced with either complete growth medium or osteogenic/adipogenic induction medium, depending on the experimental design. The sequences of the siRNA used are provided in **Supplementary Table S3**.

### Luciferase assay

A 2051-bp fragment of the *SLC3A1* promoter region (from −2000 to +50 bp) was synthesized by General Biol (Anhui) Co., Ltd. (Anhui, China) and cloned into the pGL3-Basic luciferase reporter vector (Promega, Madison, WI). Site-directed mutagenesis of the putative SMAD3 binding sites was performed using the Mut Express II Fast Mutagenesis Kit (Vazyme). The mutagenic primer sequences were as follows: Mutant site 1-F: 5′-GGCCAA<u>A</u>G<u>ACGACA</u>TCAAACTCCTGCCCTTGGTG-3′; Mutant site 1-R: 5′-TTTGA<u>TGTCGT</u>C<u>T</u>TGGCCAACATGGCCAAACCC-3′; Mutant site 2-F: 5′-<u>A</u>G<u>AC</u>G<u>ACA</u>TTTGTCCAGCTAGGTTTATTTATAAACC-3′; Mutant site 2-R: 5′-GCTGGACAAA<u>TGT</u>C<u>GT</u>C<u>T</u>AAGGAAGAAAGGGTAAAAGTACAGAGG-3′ (Mutated sequences are underlined). HEK293T cells were transfected with the luciferase constructs along with either pECMV-3×FLAG or pECMV-3×FLAG-SMAD3 using Lipofectamine 2000 (ThermoFisher Scientific). After 72 hours, cells were lysed, and luciferase activity was measured using the Dual-Luciferase Reporter Assay System (Promega). Luminescence was detected using a SpectraMax i3x Multi-Mode Microplate Reader (Molecular Devices) in luminescence mode. Firefly luciferase activity was normalized to Renilla luciferase for quantification.

### Cystine uptake assay

Cells were seeded at a density of 2 × 10^5^ cells/mL into an 8-well chamber slide (Labselect, Hefei, Anhui, China). After 24 h, the culture medium was replaced with growth medium containing FITC-labeled cystine (Ruixi Biotechnology Co., Ltd, Xi’an, Shaanxi, China). Following a 72-hour incubation, the supernatant was discarded and cells were gently washed with PBS to remove excess fluorescent substrate. Cellular uptake of FITC-cystine was visualized using a Nikon Ti2-E fluorescence microscope (Nikon, Tokyo, Japan).

### Cysteine detection

Intracellular cysteine levels were measured using a commercial cysteine assay kit (Elabscience Biotechnology Co., Ltd., Wuhan, Hubei, China). MSCs were seeded in 12-well plates at a density of 2 × 10^5^ cells/mL. After 3 days of treatment according to experimental grouping, the culture medium was removed and cells were washed with PBS. Cells were then processed following the manufacturer’s instructions. Absorbance was measured at 600 nm using a microplate reader.

### Reduced glutathione (GSH) detection

MSCs were treated under the indicated conditions for 3 days. After removal of the supernatant, cells were washed with PBS, digested, and harvested by centrifugation. Cell pellets were subjected to two rapid freeze-thaw cycles using liquid nitrogen and a 37 °C water bath. Subsequently, the samples were centrifuged at 10,000 × g for 10 minutes at 4 °C, and the supernatant was collected. Intracellular levels of reduced GSH were then quantified using a commercial GSH Detection Kit (Nanjing Jiancheng Bioengineering Institute, Nanjing, Jiangsu, China).

### Reactive oxygen species (ROS) detection

MSCs were seeded in 48-well plates at a density of 2 × 10^5^ cells/mL. Intracellular ROS levels were assessed using a ROS Assay Kit (Beyotime Biotech Inc.) based on the fluorescent probe DCFH-DA. After treatment for 3 days, the culture supernatant was removed, and cells were gently washed with PBS. Subsequently, cells were incubated with 10 μM DCFH-DA at 37 °C for 20 minutes in the dark. After incubation, cells were washed three times with PBS to eliminate excess dye. Fluorescence intensity was then measured using a SpectraMax® i3x Multi-Mode Microplate Reader (Molecular Devices).

### Enzyme-linked immunosorbent assay (ELISA)

ELISA was performed to quantify the levels of cystine in culture supernatants, cell lysates, and mouse/human serum using the kit from YaJi Biological Technology Co., Ltd (Shanghai, China) and Jiangsu Meimian Industrial Co., Ltd (Yancheng, Jiangsu, China), according to the manufacturer’s instructions. Absorbance was measured at 450 nm using a microplate reader.

### JC-1 staining for mitochondrial membrane potential

Mitochondrial membrane potential was assessed using an enhanced JC-1 detection kit (Beyotime Biotech Inc.). MSCs were incubated with JC-1 staining solution at 37 °C for 20 minutes. After staining, the supernatant was discarded and cells were washed twice with the JC-1 buffer solution. Culture medium was added to each well, and fluorescence was observed and imaged using a Nikon Ti2-E fluorescence microscope.

### Cell proliferation assay

Cell proliferation was assessed using a Cell Counting Kit-8 (CCK-8; Vazyme). MSCs (1000 cells/well) were seeded in 96-well plates. On day 3, 10 μL of CCK-8 solution was added to each well and incubated for 1 hour at 37 °C. Absorbance was measured at 450 nm using a microplate reader.

### Senescence-associated β-galactosidase (SA-β-Gal) staining

Cellular senescence was assessed using a Senescence β-Galactosidase Staining Kit (Vazyme) according to the manufacturer’s protocol. MSCs were seeded in 48-well plates at a density of 2 × 10⁴ cells/mL. After experimental treatments, the medium was removed and cells were washed with PBS. Cells were then fixed with 150 μL of fixative solution per well for 15 minutes at room temperature, followed by three washes with PBS. Subsequently, cells were incubated with staining working solution overnight at 37 °C in a CO_2_-free and dark environment. Stained cells were visualized and imaged using a standard bright-field microscope.

### Construction of stably transfected MSCs

Human adipose-derived MSCs were seeded in 6-well plates at 50-60% confluency and transduced with lentivirus encoding human SLC3A1 (OBiO Technology (Shanghai) Corp., Ltd, Shanghai, China) at a multiplicity of infection (MOI) of 50. After 24 hours, the culture medium was replaced with fresh complete medium. At 72 hours post-infection, transduced cells were selected with 1 µg/mL puromycin for 7 days to establish a stable overexpression line. Puromycin-resistant cells were expanded and used for subsequent experiments.

### Ectopic bone formation

A total of 2.5 × 10^6^ MSCs were resuspended and mixed with 45 mg of β-tricalcium phosphate (β-TCP) ceramic particles (Shanghai Bio-lu Biomaterials Co., Ltd., Shanghai, China). The cell-scaffold constructs were subcutaneously implanted into the dorsal region of 8-week-old immunodeficient (nude) mice. Mice received cystine supplementation via drinking water at a concentration of 20 mg/L. After 8 weeks, the mice were euthanized, and the implants were harvested for further histological and imaging analyses.

### Calvarial bone defects

Mice were anesthetized using 2%-3% isoflurane delivered at a flow rate of 1 L/min. Following a midline scalp incision, a full-thickness circular defect (4.0 mm in diameter) was created in the parietal bone, away from cranial sutures, using a sterile trephine drill. A total of 2 × 10^6^ MSCs suspended in hydrogel matrix were carefully implanted into the defect site. The overlying skin was sutured, and animals were monitored postoperatively. Mice received cystine supplementation via drinking water at a concentration of 20 mg/L. Eight weeks after surgery, the mice were euthanized and calvarial bones were harvested for imaging and histological evaluation.

### Tibial defect model

Mice were anesthetized using 2%-3% isoflurane at a flow rate of 1 L/min. A monocortical circular defect with a diameter of 1.0 mm was created in the proximal tibia using a dental drill under sterile conditions. The skin was then sutured with sutures. Starting on the day of surgery, mice received cystine supplementation via drinking water at a concentration of 20 mg/L, refreshed every 3 days. After 2 weeks, mice were euthanized and tibial samples were collected for subsequent analysis.

### Ovariectomy (OVX) model

Eight-week-old female mice were anesthetized with 2%-3% isoflurane delivered at a flow rate of 1 L/min. Bilateral ovariectomy was performed via dorsolateral incisions, followed by layered closure of the muscle and skin using sutures. Sham-operated mice underwent identical surgical procedures without ovary removal. After surgery, mice in the treatment group received cystine supplementation in their drinking water at a concentration of 20 mg/L, with fresh solution provided every 3 days. After 8 weeks of treatment, animals were euthanized and bone tissues were harvested for further analysis.

### Micro-computed tomography (micro-CT) and histomorphometry analyses

Bone specimens were fixed in 4% PFA at 4 °C overnight. Micro-CT imaging was performed using a Scanco Medical CT-80 scanner (Scanco Medical AG, Bruttisellen, Switzerland) at 55 kV and 144 μA, with a resolution of 12 μm. For three-dimensional morphometric analysis, image datasets were processed using CTAn, CTVol, and Dataview software (all from SkyScan, Bruker, Kontich, Belgium). Following radiographic scanning, samples were decalcified in 0.5 mM EDTA for 30-60 days, embedded in OCT compound, and cryosectioned at a thickness of 20 μm. Hematoxylin and eosin (H&E) staining and alkaline phosphatase (ALP) staining were performed on serial sections (Biosharp; Beyotime Biotech Inc.). Tartrate-resistant acid phosphatase (TRAP) staining was performed using a TRAP staining kit (Servicebio, Wuhan, Hubei, China).

### Immunofluorescence Staining

For tissue sections, samples were washed and incubated with antigen retrieval solution at RT for 3-5 minutes. Cultured cells were fixed with 4% PFA for 15 minutes, followed by permeabilization with 0.2% Triton X-100 for 10 minutes. Prior to staining, both tissue sections and cultured cells were blocked with 5% normal goat serum (Jackson Immunoresearch Laboratories, West Grove, PA) for 1 hour at RT. Primary antibodies were applied and incubated overnight at 4 °C. The next day, samples were incubated with secondary antibodies for 1.5 hours at RT. Nuclei were counterstained using SuperKine™ Enhanced Antifade Mounting Medium (Abbkine, Wuhan, Hubei, China), and samples were subsequently imaged. Antibody details are listed in **Supplementary Table S1**.

### Human specimen collection

Human serum samples were obtained from patients treated at the Department of Spine Surgery, Nanfang Hospital, Southern Medical University. Osteoporotic and non-osteoporotic subjects were enrolled according to clinical diagnostic criteria. Human bone tissue specimens were collected from osteoporotic and non-osteoporotic patients undergoing surgical treatment at Nanfang Hospital, Southern Medical University. All human sample collection procedures were approved by the Ethics Committee of Nanfang Hospital, Southern Medical University. Samples are summarized in **Supplementary Table S7**.

### Statistical Analysis

All data were presented as mean ± standard deviation (SD). Statistical significance was evaluated using either a two-tailed Student’s *t*-test or one-way/two-way ANOVA, as appropriate, with GraphPad Prism version 10.4. Differences were considered statistically significant at *P*<0.05 (*), *P*<0.01 (**), and *P*<0.001 (***).

## Data availability

The raw RNA sequencing data from this study have been deposited in the Genome Sequence Archive for Human^46,47^ in National Genomics Data Center^48^, China National Center for Bioinformation/Beijing Institute of Genomics, Chinese Academy of Sciences (GSA-human: HRA018721). These datasets are publicly accessible at https://ngdc.cncb.ac.cn/gsa-human. In addition, publicly available single-cell RNA-seq datasets were obtained from the Gene Expression Omnibus (GEO) database under accession number GSE147287.

## Declaration of generative AI and AI-assisted technologies in the writing process

During the preparation of this work, the authors used ChatGPT (OpenAI) in order to improve language expression and grammatical accuracy. After using this tool, the authors reviewed and edited the content as needed and take full responsibility for the content of the publication.

## Acknowledgments

This work was funded by National Key R&D Program of China (Nos. 2022YFC2502901, 2022YFC2502903), National Natural Science Foundation of China (No. 82472446), Guangdong Basic and Applied Basic Research Foundation (No. 2022A1515110127), and Qinghai Key Laboratory Of Tibetan Medicine Pharmacology and Safety Evaluation, Northwest Institute of Plateau Biology, Chinese Academy of Sciences (2024-ZY-01). Figure 8 were created with BioRender.com.

## Author Contributions

J.X. conceptualized the study. Y.G., Q.L., Q.X., S.C., L.J., Y.H., and J.X. conducted the experiments and analyzed the data. J.C. provided human adipose tissue specimens obtained from liposuction procedures. A.W.J. and Z.Z. assisted with data interpretation and commented on the manuscript. Y.G. and J.X. drafted the manuscript and prepared the original version of the study. Z.Z. and J.X. revised and refined the manuscript. J.X. were responsible for funding acquisition.

