## Supplementary materials for "Nutrient flux governs osteogenic fate commitment through the SLC3A1-cystine Axis"

### Supplementary Figures

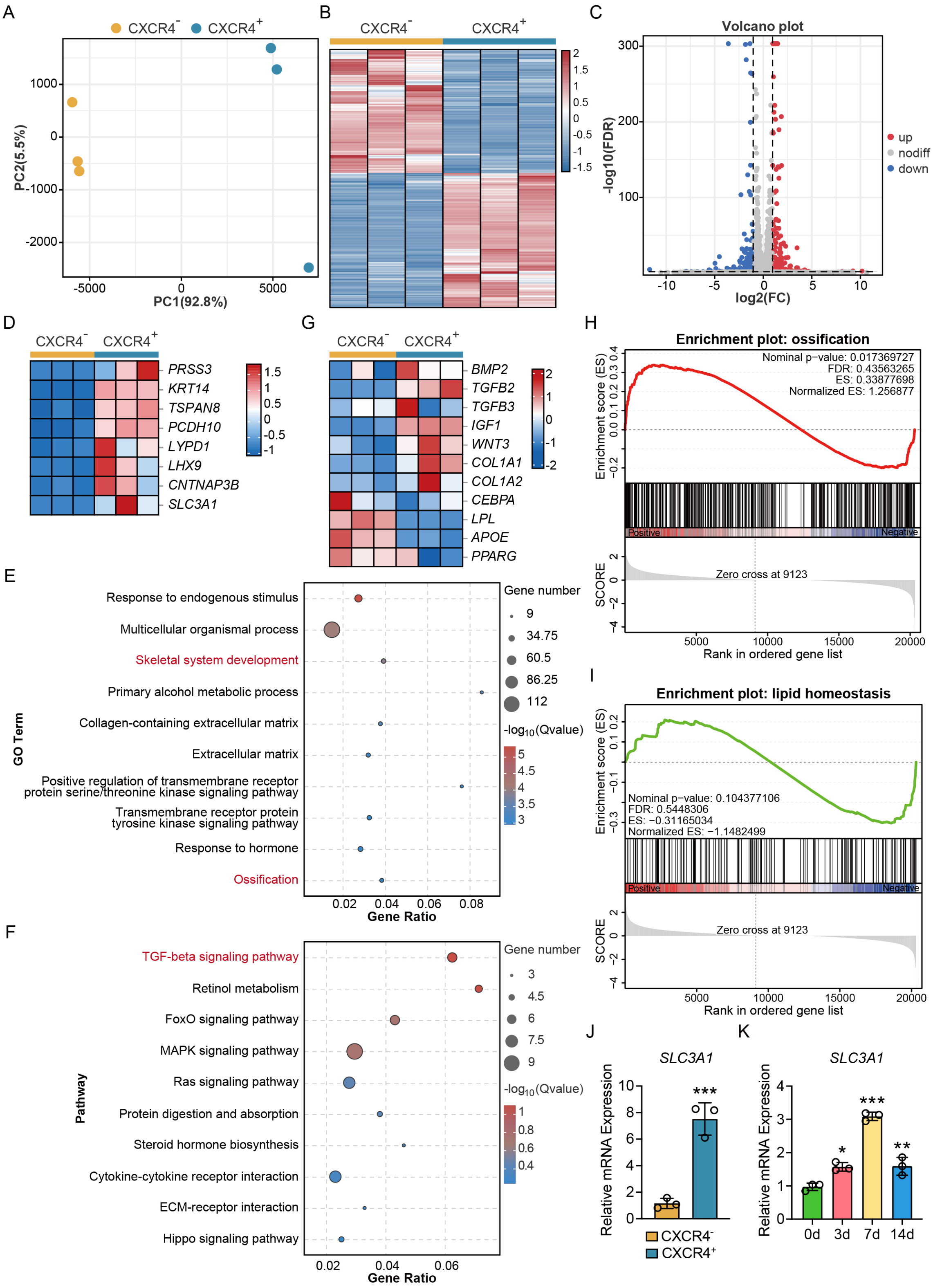


**Supplementary Figure 1. Transcriptomic profiling reveals SLC3A1 as a candidate regulator enriched in CXCR4⁺ mesenchymal cells. (A)** Principal component analysis (PCA) showing clear transcriptional separation between CXCR4⁻ and CXCR4⁺ cells based on bulk RNA-seq. **(B)** Hierarchical clustering heatmap of differentially expressed genes (DEGs) between $CXCR4^-$ and $CXCR4^+$ groups. **(C)** Volcano plot highlighting DEGs in $CXCR4^-$ and $CXCR4^+$ cells (red: upregulated in $CXCR4^+$ MSCs, blue: downregulated in $CXCR4^+$ MSCs; FDR < 0.05, $|\log_2 FC| > 1$). **(D)** Top eight genes upregulated in $CXCR4^+$ cells identified by DEG analysis. **(E)** GO enrichment analysis of DEGs showing upregulation of biological processes related to skeletal development and ossification. **(F)** KEGG pathway analysis demonstrating enrichment of TGF-β signaling, MAPK signaling, and cytokine-cytokine receptor interaction in $CXCR4^+$ cells. **(G)** Expression heatmap of lineage-specific markers showing osteogenic bias in $CXCR4^+$ cells and adipogenic bias in $CXCR4^-$ cells. **(H,I)** GSEA indicated significant enrichment of ossification-related gene sets and downregulation of lipid homeostasis pathway in $CXCR4^+$ cells. **(J)** qPCR validation of top DEG candidates, confirming that SLC3A1 was selectively elevated in $CXCR4^+$ cells. Other genes showed inconsistent or nonsignificant differences (see **Supplementary Fig. 2**). **(K)** Temporal expression of SLC3A1 during osteogenic induction of adipose-derived MSCs. In panel K, asterisks indicate statistical significance compared with day 0. Data are presented as mean ± SD. *$P$<0.05; **$P$<0.01; ***$P$<0.001.

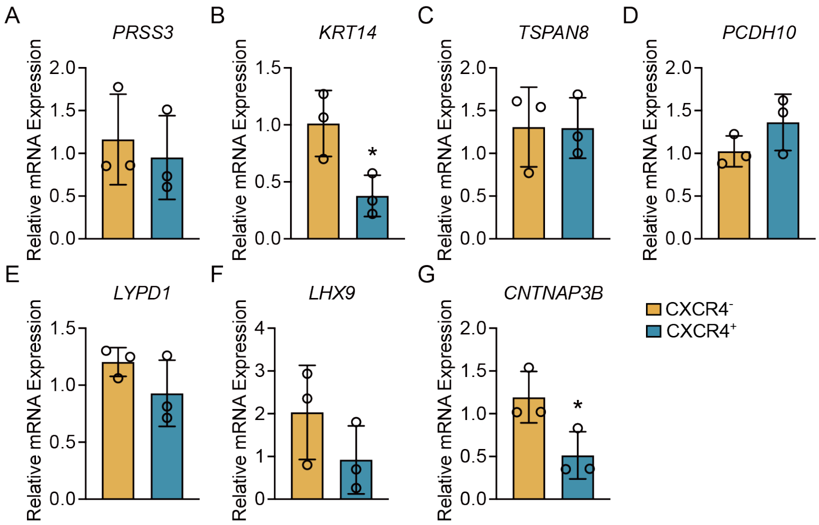


**Supplementary Figure 2.** The mRNA expression levels of selected differentially expressed candidate genes in CXCR4$^-$ and CXCR4$^+$ mesenchymal cells, as assessed by qPCR, including **(A)** *PRSS3*, **(B)** *KRT14*, **(C)** *TSPAN8*, **(D)** *PCDH10*, **(E)** *LYPD1*, **(F)** *LHX9*, and **(G)** *CNTNAP3B*. Data are presented as mean ± SD. *$P$<0.05.

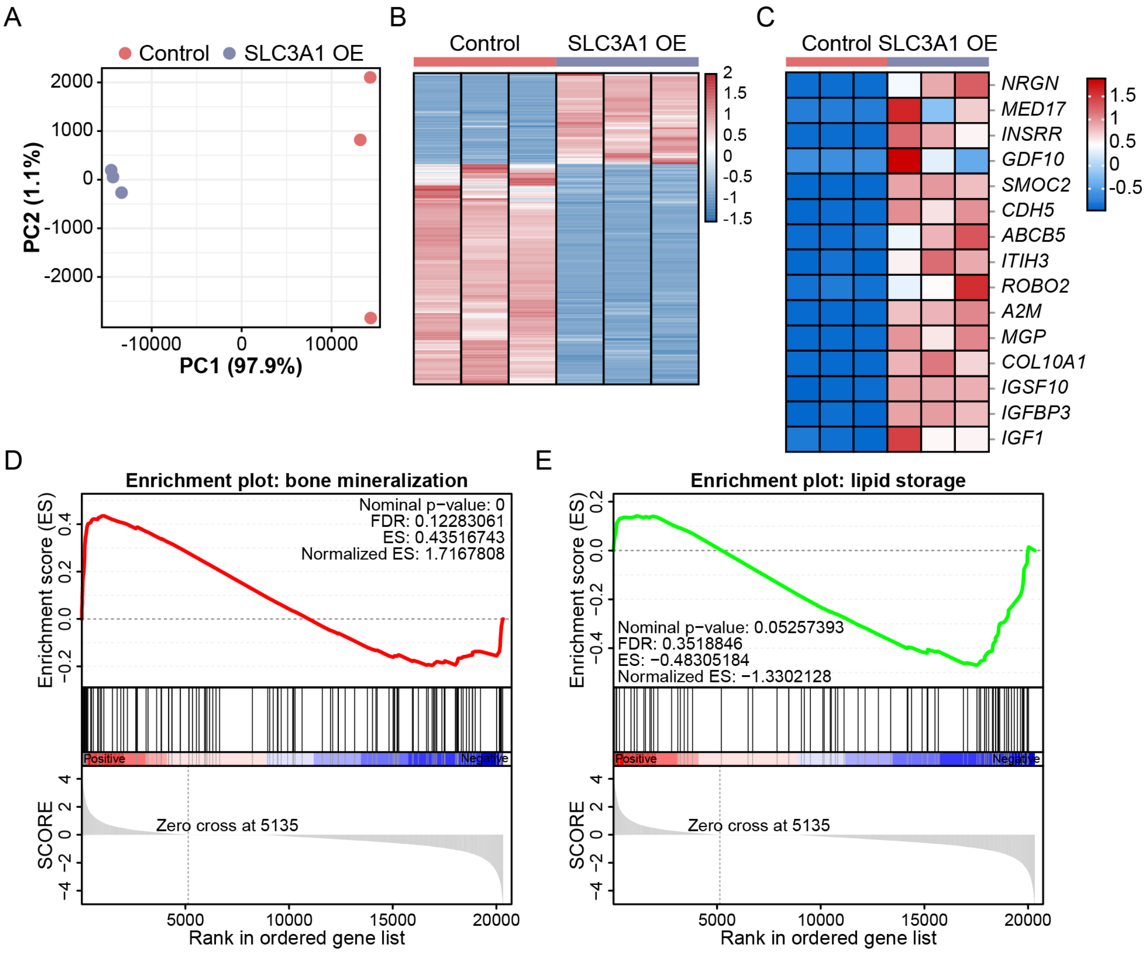


**Supplementary Figure 3. Bulk RNA sequencing among control and SLC3A1-overexpressing MSCs. (A,B)** Principal component analysis (PCA) and hierarchical clustering heatmap of bulk RNA-seq data showing distinct transcriptomic profiles between control and SLC3A1-overexpressing cells. **(C)** Top upregulated DEGs in SLC3A1-overexpressing cells. **(D)** GSEA showing activation of bone mineralization in SLC3A1 OE group. **(E)** GSEA revealing suppression of lipid storage pathway in SLC3A1 OE group. OE, overexpression.

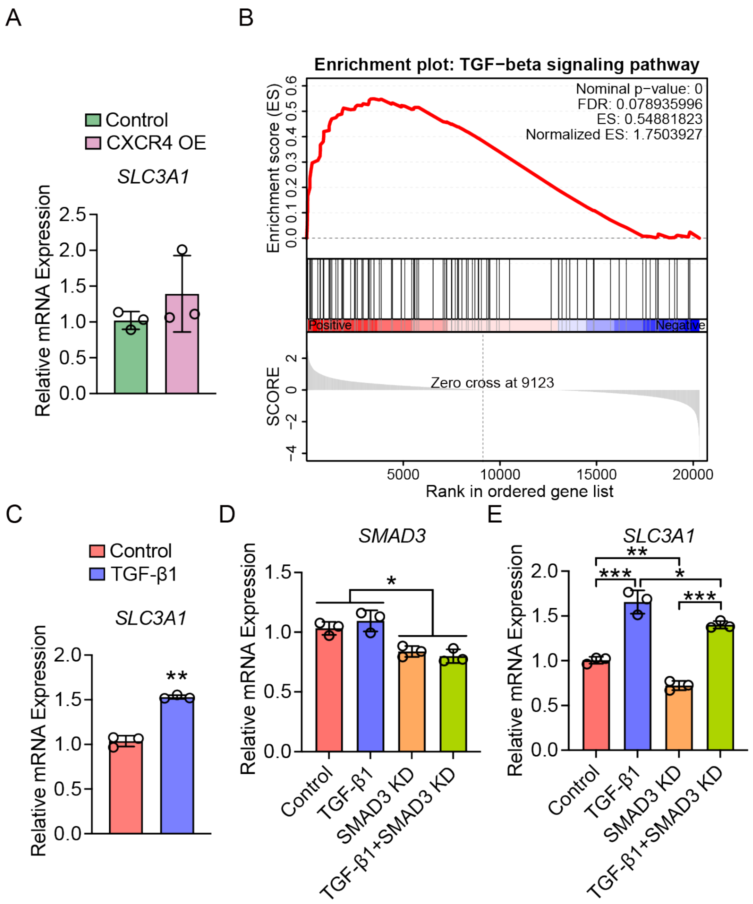


**Supplementary Figure 4. TGF-β/SMAD3 signaling upregulates SLC3A1 expression. (A)** qPCR analysis showing that overexpression of CXCR4 did not affect SLC3A1 expression. **(B)** GSEA revealed activation of the TGF-β signaling pathway in $CXCR4^{+}$ mesenchymal cells. **(C)** Stimulation with TGF-β1 (2 ng/mL) significantly increased *SLC3A1* mRNA levels in MSCs. **(D)** qPCR validation of *SMAD3* knockdown efficiency using siRNA. **(E)** *SMAD3* knockdown suppressed TGF-β1-induced upregulation of SLC3A1 expression, indicating that SMAD3 mediates the TGF-β-SLC3A1 regulatory axis. OE, overexpression; KD, knockdown. Data are presented as mean ± SD. **P*<0.05; ***P*<0.01; ****P*<0.001.

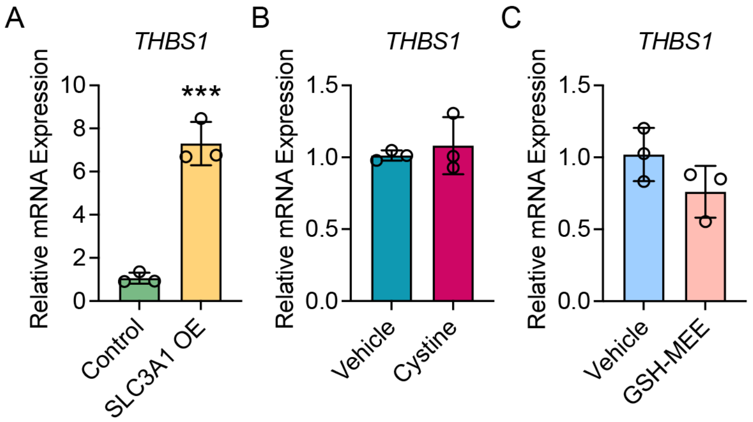


**Supplementary Figure 5. qPCR analysis of *THBS1* expression under different treatment conditions.** **(A)** Relative *THBS1* expression following SLC3A1 overexpression. **(B)** Relative *THBS1* expression after cystine treatment. **(C)** Relative *THBS1* expression after treatment with glutathione monoethyl ester (GSH-MEE). OE, overexpression. Data are presented as mean ± SD. ***$P$<0.001.

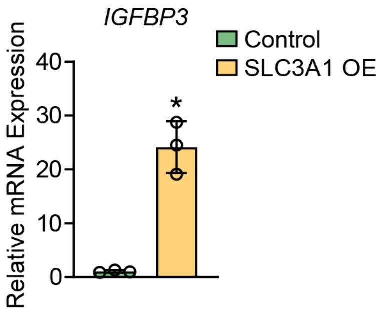


**Supplementary Figure 6. qPCR analysis of *IGFBP3* expression following SLC3A1 overexpression.** OE, overexpression. Data are presented as mean ± SD. *$P<0.05$.

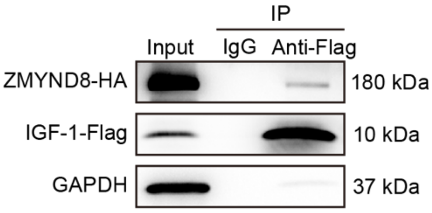


**Supplementary Figure 7. Co-immunoprecipitation (Co-IP) confirms the interaction between IGF-1 and ZMYND8.** HEK293T cells were transfected with IGF-1-Flag and ZMYND8-HA expression plasmids. Cell lysates were subjected to immunoprecipitation using an anti-Flag antibody. Input, IgG control, and anti-Flag IP groups were analyzed by western blot. Blots were probed with antibodies against HA, Flag, and GAPDH. ZMYND8 was detected specifically in the anti-Flag immunoprecipitates, indicating its interaction with IGF-1.

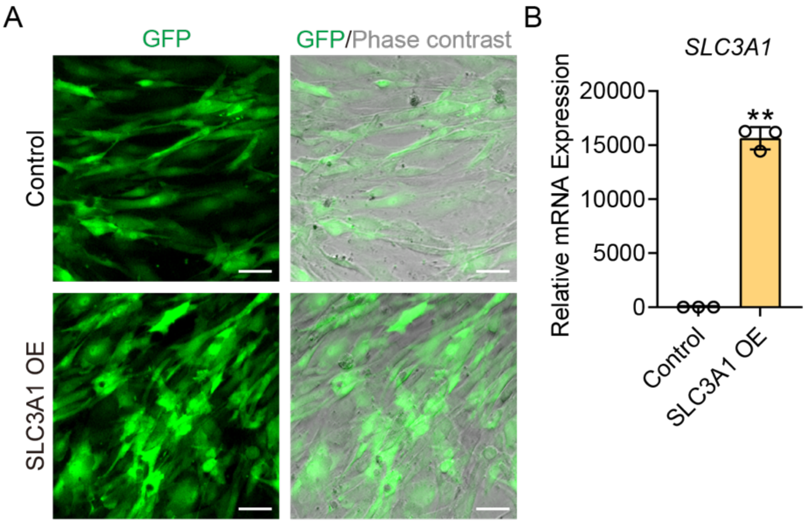


**Supplementary Figure 8. Stable overexpression of SLC3A1 in MSCs and validation of transfection efficiency. (A)** Adipose-derived MSCs were stably transfected with control GFP lentivirus or SLC3A1-overexpressing lentivirus (SLC3A1 OE). GFP fluorescence and phase contrast images were captured 10 days post-transduction to confirm successful lentiviral infection. Scale bars: 50 µm. **(B)** Overexpression efficiency of *SLC3A1* in stably transfected cells was assessed by qPCR following puromycin selection. OE, overexpression. Data are presented as mean ± SD. $^{**}P<0.01$.

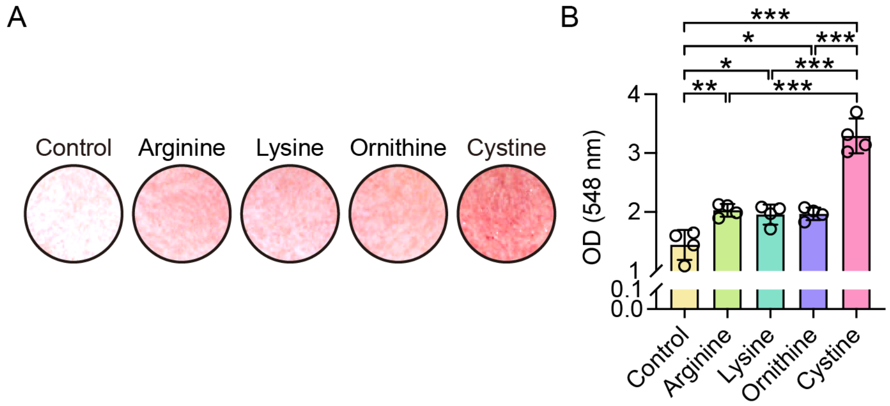


**Supplementary Figure 9. Cystine preferentially enhances osteogenic differentiation relative to other dibasic amino acids. (A,B)** Alizarin Red S staining **(A)** and quantitative analysis **(B)** showing osteogenic mineralization following treatment with cystine or other dibasic amino acids, including arginine, lysine, and ornithine, all at a concentration of 1 mM. Data are presented as mean ± SD. **P*<0.05; ***P*<0.01; ****P*<0.001.

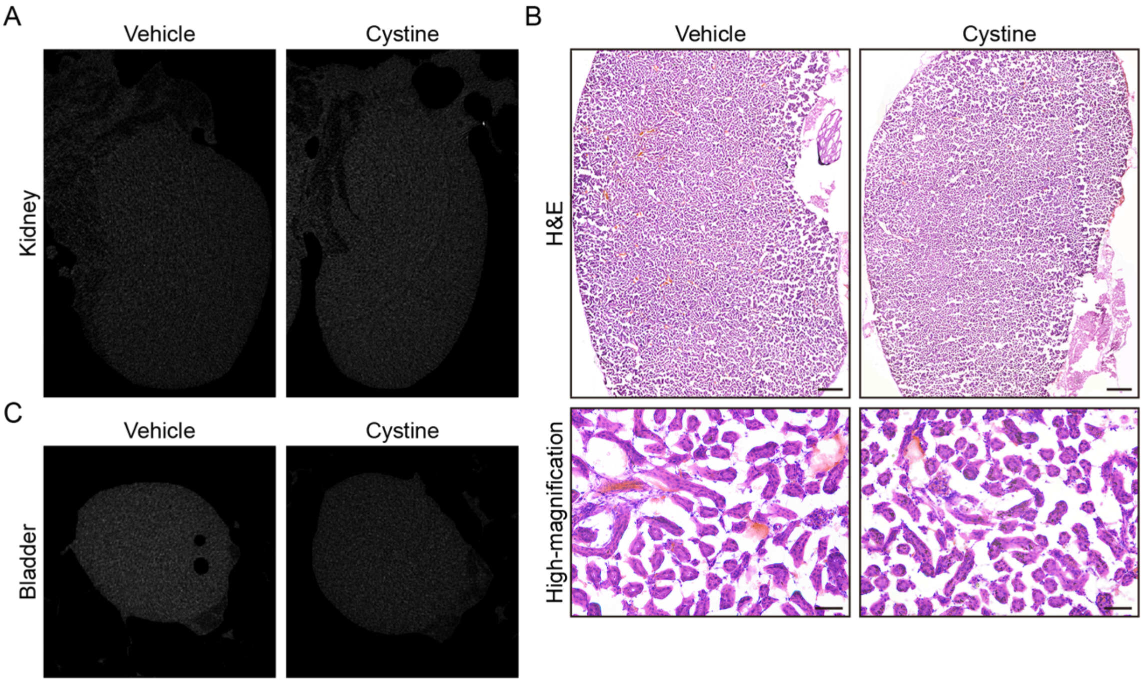


**Supplementary Figure 10. Evaluation of renal and bladder safety following oral cystine supplementation. (A)** Representative micro-CT images of kidneys from mice receiving cystine-supplemented drinking water, showing no evidence of ectopic calcification or kidney stone formation. **(B)** H&E staining of kidney sections revealed no histopathological abnormalities, indicating preserved renal architecture and cystine safety. Scale bars: 200 µm (upper panel) and 50 µm (lower panel). **(C)** Micro-CT analysis of bladder tissue also showed absence of calcified signals, ruling out cystine-induced bladder stone formation.

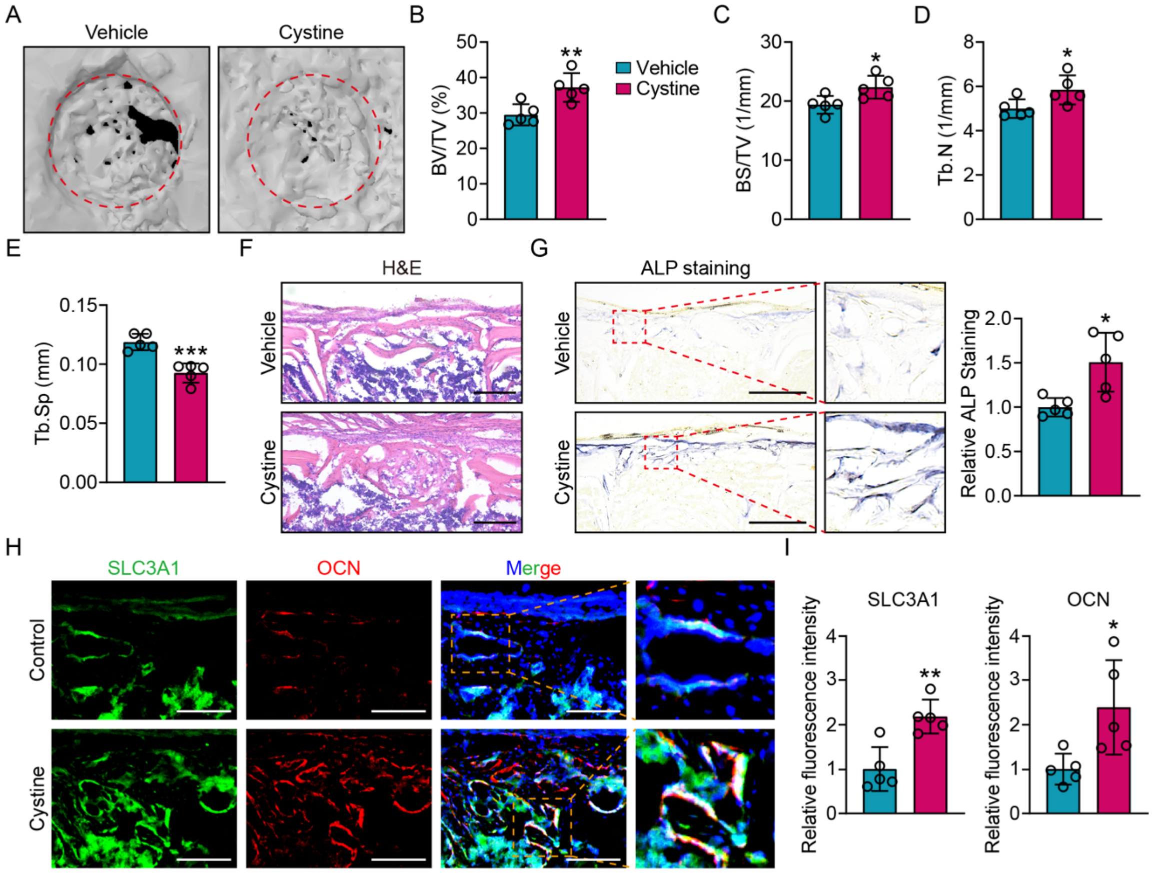


**Supplementary Figure 11. Cystine supplementation accelerates tibial defect healing *in vivo*. (A)** Representative micro-CT images of monocortical tibial defects 14 days post-surgery in vehicle- and cystine-treated groups. Red circles mark the boundaries of the original defect sites. **(B-E)** Quantitative micro-CT analysis showing increased **(B)** BV/TV, **(C)** BS/TV, **(D)** Tb.N, and decreased **(E)** Tb.Sp in the cystine group, indicating enhanced bone regeneration. **(F)** H&E staining of defect sites revealed more robust bone formation in cystine-treated mice. **(G)** Representative and quantitative analysis of ALP staining demonstrated elevated osteogenic activity following cystine supplementation. **(H)** Immunofluorescence co-staining for SLC3A1 and OCN in defect regions showed increased expression and co-localization in the cystine group. Nuclei were counterstained with DAPI (blue). **(I)** Quantification of fluorescence intensity of SLC3A1 and OCN revealed significantly elevated levels in cystine-treated samples compared to vehicle controls. *n* = 5 per group. Scale bars: 200 μm (**F,G**) and 100 μm (**H**). Data are presented as mean ± SD. *$P$<0.05; **$P$<0.01; ***$P$<0.001.

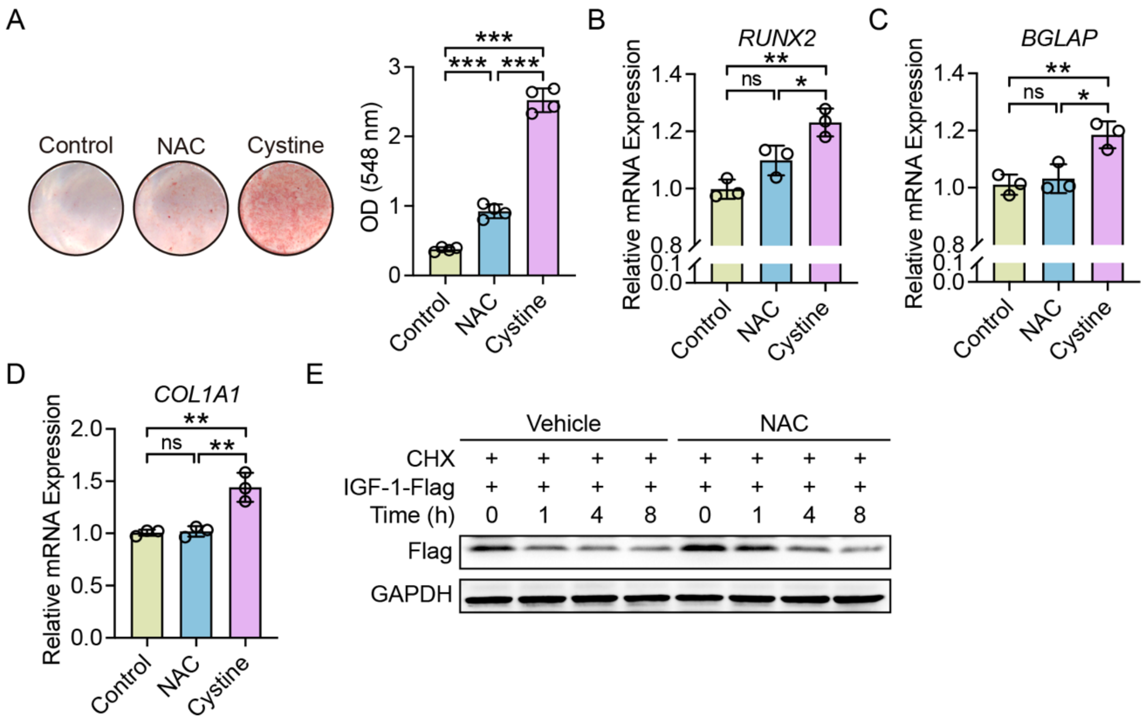


**Supplementary Figure 12. Comparative effects of N-acetylcysteine (NAC) and cystine on osteogenic differentiation. (A)** Representative Alizarin Red S staining images of adipose-derived MSCs treated with control, NAC (1 mM), or cystine (1 mM) following 9 days of osteogenic induction. **(B-D)** qPCR analysis of osteogenic marker genes *RUNX2* **(B)**, *BGLAP* **(C)**, and *COL1A1* **(D)**. Gene expression levels were normalized to GAPDH and are presented relative to the control group. **(E)** NAC treatment did not alter the degradation kinetics of IGF-1 protein. HEK293T cells were treated with either vehicle or NAC, followed by CHX treatment for the indicated time periods. Protein levels of IGF-1-Flag were detected by western blot. Data are presented as mean ± SD. ns, not significant. *$P$<0.05; **$P$<0.01; ***$P$<0.001.

**Supplementary Tables**

**Table S1. Antibodies used.**

| Antibody | Company | Catalog # | Use |
|---|---|---|---|
| Rabbit anti-Human ALP | HuaBio | ET1601-21 | WB |
| Rabbit anti-Human α-SMA | Affinity | AF1032 | IF |
| Mouse anti-Human CXCR4-APC | BioLegend | 306510 | FC |
| Mouse anti-Human CXCR4 | R&D | MAB170 | Neutralization |
| Mouse anti-Flag Tag (HRP Conjugated) | Beyotime | AF2855 | WB |
| Rabbit anti-Mouse GAPDH | HuaBio | ET1601-4 | WB |
| Rabbit anti-Mouse/Human IGF-1 | MedChemExpress | HY-P81112 | IF |
| Mouse anti-HA Tag (HRP Conjugated) | Beyotime | AF2861 | WB |
| Rabbit anti-Mouse Osteocalcin | Affinity | DF-12303 | IF |
| Rabbit anti-Human $p16^{INK4A}$ | HuaBio | ER1803-53 | IF |
| Rabbit anti-Human RUNX2 | HuaBio | ET1612-47 | WB |
| Mouse anti-Human SLC3A1 | Santa Cruz Biotechnology | sc-518240 | IF, FC |
| Rabbit anti-Human phospho-SMAD3 | Cell Signaling Technology | 9520 | CUT&Tag |
| Mouse anti-Ubiquitin | Abmart | M026378 | WB |
| Goat anti-Mouse AF594 | Abcam | ab150116 | IF |
| Goat anti-Rabbit AF488 | Abcam | ab150077 | IF |
| Goat anti-Rabbit iFluor™ 594 | HuaBio | HA1122 | IF |
| Goat anti-Mouse iFluor™ 647 | HuaBio | HA1127 | IF |
| HRP-labeled Goat anti-Rabbit IgG(H+L) | Biodragon | BF03008X | WB |
| HRP Conjugated Goat anti-Mouse IgG | HuaBio | HA1006 | WB |
| HRP Conjugated Anti-mouse IgG for IP Nano-secondary antibody | HuaBio | NBI02H | WB |

CUT&Tag: Cleavage Under Targets and Tagmentation; FC: Flow cytometry; IF: Immunofluorescent staining; WB: Western blot.

### Table S2. qPCR primers used.

| Genes (Human) | Forward | Reverse |
|---|---|---|
| *ALP* | 5’-ACCACCACGAGAGTGAACCA-3’ | 5’-CGTTGTCTGAGTACCAGTCCC-3’ |
| *BGLAP* | 5’-CGCTACCTGTATCAATGGCTGG-3’ | 5’-CTCCTGAAAGCCGATGTGGTCA-3’ |
| *CEBPA* | 5’-AGGAGGATGAAGCCAAGCAGCT-3’ | 5’-AGTGCGCGATCTGGAACTGCAG-3’ |
| *CNTNAP3B* | 5’-ACCTTTCAGTTATCCTCGCCA-3’ | 5’-AATATGAGAGATTTGACGGCGTT-3’ |
| *CDKN1A* | 5’-CGATGGAACTTCGACTTTGTCA-3’ | 5’-GCACAAGGGTACAAGACAGTG-3’ |
| *CDKN2A* | 5’-GGGTTTTCGTGGTTCACATCC-3’ | 5’-CTAGACGCTGGCTCCTCAGTA-3’ |
| *COL1A1* | 5’-GAGGGCCAAGACGAAGACATC-3’ | 5’-CAGATCACGTCATCGCACAAC-3’ |
| *FABP4* | 5’-ACTGGGCCAGGAATTTGACG-3’ | 5’-CTCGTGGAAGTGACGCCTT-3’ |
| *GAPDH* | 5’-CTGGGCTACACTGAGCACC-3’ | 5’-AAGTGGTCGTTGAGGGCAATG-3’ |
| *IGF1* | 5’-CTCTTCAGTTCGTGTGTGGAGAC-3’ | 5’-CAGCCTCCTTAGATCACAGCTC-3’ |
| *IGFBP3* | 5’-AGACACACTGAATCACCTGAAGT-3’ | 5’-AGGGCGACACTGCTTTTTCTT-3’ |
| *KRT14* | 5’-TGAGCCGCATTCTGAACGAG-3’ | 5’-GATGACTGCGATCCAGAGGA-3’ |
| *LHX9* | 5’-GATGGAGCGCAGATCCAAGAC-3’ | 5’-CCAGCAGATAGTACCTGTCCG-3’ |
| *LPL* | 5’-TCATTCCCGGAGTAGCAGAGT-3’ | 5’-GGCCACAAGTTTTGGCACC-3’ |
| *LYPD1* | 5’-GGCAACTTTTTGCGGATTGTT-3’ | 5’-CGTTCACCGTGCAATTCACA-3’ |
| *PCDH10* | 5’-TGGATGGTGGAAGGAGTCTTT-3’ | 5’-TTCAGCGATATTCCCCACGAA-3’ |
| *PPARG* | 5’-GGGATCAGCTCCGTGGATCT-3’ | 5’-TGCACTTTGGTACTCTTGAAGTT-3’ |
| *PRSS3* | 5’-GCTGAGTGTAAAGCCTCCTACC-3’ | 5’-CAACTCCTTGGAGCTGTCCGTT-3’ |
| *RUNX2* | 5’-TGGTTACTGTCATGGCGGGTA-3’ | 5’-TCTCAGATCGTTGAACCTTGCTA-3’ |
| *SMAD3* | 5’-TGGACGCAGGTTCTCCAAAC-3’ | 5’-CCGGCTCGCAGTAGGTAAC-3’ |
| *SLC3A1* | 5’-CAGGAGCCCGACTTCAAGG-3’ | 5’-GAGGGCAATGATGGCTATGGT-3’ |
| *SP7* | 5’-CCTCTGCGGGACTCAACAAC-3’ | 5’-AGCCCATTAGTGCTTGTAAAGG-3’ |
| *Spike-in* | 5’-GCCTTCTTCCCATTTCTGATCC-3’ | 5’-CACGAATCAGCGGTAAAGGT-3’ |
| *THBS1* | 5’-TCTGCAACAAGCAGGACTGT-3’ | 5’-CCATTGTGGTTGAAGCAGGC-3’ |
| *TSPAN8* | 5’-ACTTCTTGTTCTGGCTATGTGG-3’ | 5’-CACAGCAACGTAGGAGCTAGA-3’ |
| *TP53* | 5’-GAGGTTGGCTCTGACTGTACC-3’ | 5’-TCCGTCCCAGTAGATTACCAC-3’ |
| CUT&Tag Site 1 | 5’-ACCACGCCTGGCTAATTTTT-3’ | 5’-GCCTGTAATCCCAGCACTTTG-3’ |
| CUT&Tag Site 2 | 5’-CCTTTCTTCCTTGGCTGGACT-3’ | 5’-GTGGAAGAGTGGCTTGCTGA-3’ |

**Table S3. siRNA sequences used in gene knockdown.**

| Genes | Target (sense) sequence (5' to 3') |
|---|---|
| Negative control | TTCTCCGAACGTGTCACGT |
| *IGF1* siRNA | CACAAATGCATGGGTGTTGTA |
| *SLC3A1* siRNA | GCCATACATGATAAAGGTTTA |
| *SMAD3* siRNA | GCCTCAGTGACAGCGCTATTT |

**Table S4. List of genes enriched in the KEGG focal adhesion pathway.**

| Focal adhesion | | | | | |
|---|---|---|---|---|---|
| *PDGFRA* | *VAV1* | *EGF* | *PIK3R2* | *VTN* | *BIRC2* |
| *FLNC* | *ITGA11* | *MYLK3* | *IGF1* | *LAMB3* | *PIK3CA* |
| *ITGB1* | *PAK5* | *VAV2* | *RASGRF1* | *IBSP* | *JUN* |
| *PARVA* | *ITGA1* | *PPP1R12C* | *ARHGAP5* | *LAMA2* | *RAC1* |
| *ITGA6* | *COL4A5* | *FLT4* | *PIP5K1A* | *ILK* | *PAK4* |
| *VCL* | *VEGFD* | *HGF* | *MAPK10* | *TLN2* | *ACTG1* |
| *PIK3CD* | *ITGA2* | *TLN1* | *BRAF* | *PIP5K1C* | *ITGA7* |
| *MAPK8* | *COL9A3* | *AKT2* | *LAMC1* | *PTK2* | *VASP* |
| *PPP1CA* | *XIAP* | *BCAR1* | *TNR* | *RELN* | *ITGA4* |
| *FLT1* | *PGF* | *RAP1B* | *PDPK1* | *THBS3* | *ITGB5* |
| *ROCK1* | *LAMA1* | *ITGB3* | *THBS2* | *ZYX* | *FLNB* |
| *COL4A2* | *COMP* | *CTNNB1* | *CCND2* | *CCND2* | *CCND1* |
| *COL4A1* | *PPP1R12A* | *CHAD* | *ACTB* | *PRKCG* | *COL4A3* |
| *COL6A2* | *SHC3* | *SOS2* | *LAMC2* | *SPP1* | *COL4A4* |
| *PDGFB* | *ITGB7* | *CAV2* | *PPP1CA* | *VEGFA* | *COL1A1* |
| *PARVG* | *ITGA5* | *LAMB4* | *LAMB1* | *DIAPH1* | *COL6A3* |
| *LAMA5* | *CRK* | *ITGAV* | *COL2A1* | *SHC2* | *PXN* |
| *ITGA8* | *FLNA* | *ITGA3* | *MYL7* | *EGFR* | *ITGA2B* |
| *RAC2* | *PPP1CB* | *DOCK1* | *BAD* | *ITGB4* | *AKT3* |
| *THBS1* | *PIP5K1B* | *MYL2* | *PIK3CB* | *PRKCB* | *MYLK* |
| *PAK6* | *THBS4* | *PRKCA* | *COL9A1* | *PAK2* | *COL6A5* |
| *MAPK1* | *CRKL* | *VEGFC* | *VWF* | *LAMB2* | *ROCK2* |
| *MYLK4* | *LAMA4* | *COL4A6* | *CAV1* | *GSK3B* | *COL6A6* |
| *TNXB* | *CAV3* | *AKT1* | *MET* | *PPP1CC* | *RAF1* |
| *SHC4* | *ITGB8* | *ITGB6* | *CCND3* | *COL9A2* | *SOS1* |
| *BCL2* | *BIRC3* | *PAK1* | *MAP2K1* | *MYL5* | *SHC1* |
| *MYL12A* | *ITGA9* | *PDGFRB* | *PPP1R12B* | *MYLPF* | *ITGA10* |
| *MYL12B* | *PAK3* | *MYL10* | *PTEN* | *COL1A2* | *RAC3* |
| *COL6A1* | *SRC* | *ACTN1* | *CAPN2* | *HRAS* | *TNN* |
| *ELK1* | *IGF1R* | *PDGFC* | *MAPK9* | *FN1* | *RHOC* |
| *PARVB* | *PIK3R1* | *ARHGAP35* | *RAPGEF1* | *GRB2* | *PIK3R3* |
| *MYL9* | *PDGFD* | *VAV3* | *VEGFB* | *RHOA* | *P3R3URF-PIK3R3* |
| *LAMA3* | *TNC* | *MAPK3* | *KDR* | *PDGFA* | *RAP1A* |
| *MYLK2* | *FYN* | *LAMC3* | *ACTN4* | *ERBB2* | *CDC42* |

**Table S5. List of genes enriched in the KEGG p53 signaling pathway.**

| **p53 signaling pathway** | | | | | |
|---|---|---|---|---|---|
| *SESN1* | *RRM2B* | *CCNB2* | *SERPINE1* | *CASP3* | *TSC2* |
| *PPM1D* | *CCND3* | *BID* | *CASP9* | *ADGRB1* | *CDK2* |
| *TP53* | *STEAP3* | *TP53I3* | *CCNG2* | *PTEN* | *ZMAT3* |
| *BBC3* | *CCND2* | *GADD45G* | *GADD45A* | *COP1* | *ATM* |
| *FAS* | *SHISA5* | *SIVA1* | *CDK4* | *CDK1* | *CCND1* |
| *AIFM2* | *DDB2* | *PERP* | *IGF1* | *TP53AIP1* | *GORAB* |
| *CDKN1A* | *CHEK2* | *CCNE1* | *CCNE2* | *IGFBP3* | *SFN* |
| *CASP8* | *MDM2* | *SIAH1* | *CDKN2A* | *RCHY1* | *MDM4* |
| *BCL2L1* | *PMAIP1* | *SESN3* | *RPRM* | *RRM2* | |
| *BAX* | *GTSE1* | *TP73* | *CDK6* | *CCNG1* | |
| *TNFRSF10B* | *THBS1* | *SESN2* | *GADD45B* | *CCNB1* | |
| *CD82* | *SERPINB5* | *CYCS* | *ATR* | *CHEK1* | |
| *PIDD1* | *BCL2* | *EI24* | *TNFRSF10A* | *APAF1* | |

**Table S6. Patient information of human specimens used for cell isolation.**

| Sample No. | Age | Gender | Tissue | Location |
|---|---|---|---|---|
| 1 | 50 | Female | Adipose | Lower back |
| 2 | 59 | Female | Adipose | Back |
| 3 | 31 | Female | Adipose | Abdomen |
| 4 | 32 | Female | Adipose | Hips |
| 5 | 29 | Female | Adipose | Abdomen |
| 6 | 25 | Female | Adipose | Abdomen |
| 7 | 49 | Female | Adipose | Posterior neck |
| 8 | 49 | Male | Bone marrow | Acetabulum |

**Table S7. Patient information of human subjects used for bone immunostaining.**

| Sample No. | Age | Gender | Group | Tissue | Location |
|---|---|---|---|---|---|
| 1 | 60 | Male | Control | Trabecular bone | Tibia |
| 2 | 82 | Male | Osteoporosis | Trabecular bone | Femoral head |

**Table S8. Patient information of human subjects used for peripheral blood serum analysis.**

| Sample No. | Age | Gender | Group | T-score |
|---|---|---|---|---|
| 1 | 63 | Female | Control | -2.3 |
| 2 | 52 | Female | | -0.2 |
| 3 | 71 | Female | | -1.5 |
| 4 | 36 | Female | | -0.9 |
| 5 | 59 | Female | | -1.4 |
| 6 | 81 | Female | | -2.1 |
| 7 | 70 | Female | | -2.3 |
| 8 | 66 | Female | | -2.3 |
| 9 | 56 | Female | | -0.7 |
| 10 | 42 | Female | | -1.8 |
| 11 | 52 | Female | | -1.1 |
| 12 | 59 | Female | | -2.3 |
| 13 | 58 | Female | | -1.6 |
| 14 | 58 | Female | | -2.1 |
| 15 | 68 | Female | | -2.0 |
| 16 | 41 | Female | | -0.3 |
| 17 | 55 | Female | | -1.8 |
| 18 | 32 | Female | | -0.8 |
| 19 | 58 | Female | | -0.9 |
| 20 | 61 | Female | | -1.8 |
| 21 | 53 | Female | Osteoporosis | -3.0 |
| 22 | 73 | Female | | -3.0 |
| 23 | 51 | Female | | -3.9 |
| 24 | 53 | Female | | -2.7 |
| 25 | 73 | Female | | -2.8 |
| 26 | 78 | Female | | -3.2 |
| 27 | 75 | Female | | -3.4 |
| 28 | 70 | Female | | -4.5 |
| 29 | 73 | Female | | -3.8 |
| 30 | 78 | Female | | -4.3 |
| 31 | 82 | Female | | -3.0 |
| 32 | 65 | Female | | -3.2 |
| 33 | 74 | Female | | -3.7 |
| 34 | 59 | Female | | -2.7 |
| 35 | 77 | Female | | -5.9 |
| 36 | 82 | Female | | -3.1 |
| 37 | 76 | Female | | -3.0 |
| 38 | 68 | Female | | -4.6 |
| 39 | 59 | Female | | -3.3 |
| 40 | 66 | Female | | -3.9 |
